# Altered TFEB subcellular localization in nigral dopaminergic neurons of subjects with prodromal, sporadic and *GBA*-related Parkinson’s disease and Dementia with Lewy bodies

**DOI:** 10.1101/2023.07.16.549189

**Authors:** Tim E Moors, Martino L Morella, Cesc Bertran-Cobo, Hanneke Geut, Vinod Udayar, Evelien Timmermans-Huisman, Angela MT Ingrassia, John JP Brevé, John GJM Bol, Vincenzo Bonifati, Ravi Jagasia, Wilma DJ van de Berg

**Author notes:** Corresponding author: Wilma D.J. van de Berg, PhD Dept. of Anatomy & Neurosciences, chair section Clinical Neuroanatomy and Biobanking Amsterdam Neuroscience Amsterdam UMC, location VU University Medical Center O2 building, room 13 E11 De Boelelaan 1108 1081 HZ Amsterdam, The Netherlands.nr.

## Abstract

Transcription factor EB is a master regulator of genes involved in the maintenance of autophagic and lysosomal homeostasis, processes which have been implicated in the pathogenesis of *GBA*-related and sporadic Parkinson’s disease (PD) and dementia with Lewy bodies (DLB). TFEB activation at the lysosomal level results in its translocation from the cytosol to the nucleus. Here, we aimed at investigating whether TFEB subcellular localization is altered in post-mortem human brain of aged individuals with either prodromal PD/DLB (incidental Lewy body disease, iLBD, N=3), *GBA*-related PD/DLB (N=9) or sPD/DLB (N=9), compared to control subjects (N=12). We scanned nigral dopaminergic neurons using high-resolution confocal and stimulated emission depletion (STED) microscopy and semi-quantitatively scored the observed TFEB subcellular localization patterns. In line with previous studies, we observed reduced nuclear TFEB immunoreactivity in PD/DLB patients compared to controls, both sporadic and *GBA*-related cases, as well as in iLBD cases. Nuclear depletion of TFEB was more pronounced in neurons with Ser129-phosphorylated (pSer129) aSyn cytopathology and in cases carrying pathogenic *GBA* variants. Interestingly, we further observed previously unidentified TFEB-immunopositive somatic clusters in human brain dopaminergic neurons and in human embryonic stem cell (hESC)-derived neurons, which localized at the Golgi apparatus. The TFEB clustering was more frequently observed and more severe in iLBD, sPD/DLB and *GBA*-PD/DLB compared to controls, particularly in pSer129 aSyn-positive neurons but also in neurons without apparent cytopathology. Notably, increased frequency of cytoplasmic TFEB clusters in aSyn-negative cells correlated with reduced total *GBA* enzymatic activity and higher Braak LB stage. In the studied patient population, altered TFEB distribution was accompanied by a reduction in overall mRNA expression levels of selected CLEAR genes, indicating a possible early dysfunction of lysosomal regulation. Overall, these findings suggest the early cytoplasmic TFEB retention and accumulation at the Golgi prior pSer129 aSyn accumulation in incidental, *GBA*-related and sporadic PD/DLB and indicate TFEB as potential as early therapeutic target for synucleinopathies

## Introduction

Parkinson’s disease (PD) and dementia with Lewy bodies (DLB) are neurodegenerative diseases pathologically defined by the presence of neuronal cytoplasmic and axonal inclusions, termed Lewy bodies (LBs) and Lewy neurites, in circumscribed regions of the brain [1]. LBs consist of a variety of membranous deposits and proteins but are mainly defined by the presence of accumulated and post-translationally modified aSyn, most notably Ser129-phosphorylated aSyn (pSer129 aSyn) [2-5]. Under physiological conditions, aSyn turnover is regulated by intracellular degradation systems, such as the ubiquitin-proteasomal system and the autophagy-lysosomal pathway (ALP) [6, 7]. In contrast, pathological forms of aSyn are not adequately degraded by these protein degradation mechanisms, leading to its accumulation in PD and DLB. Failure of the ALP has been observed in PD/DLB, a finding which is supported by GWAS and linkage studies that have identified many genetic risk factors among ALP-related genes [8-10].

The most common genetic risk factors associated with PD and DLB are heterozygous mutations in the *GBA* gene. *GBA* encodes for the enzyme ß-glucocerebrosidase (GCase), a lysosomal enzyme that catalyzes the hydrolysis of the sphingolipid glucosylceramide into glucose and ceramide [11-13]. Results of pathological and experimental studies have suggested a reciprocal relationship between GCase and aSyn [14-22], although the nature of this relationship is currently unclear. While the ability of GCase and aSyn to physically interact has been demonstrated *in vitro* using nuclear magnetic resonance spectroscopy [23, 24], alternative hypotheses propose the direct interaction between aSyn and GCase as a result of increased GCase substrate levels [14, 25-27], decreased intracellular trafficking of GCase [20, 21, 27], and/or ALP dysfunction [20, 21, 28, 29]. Indeed, induced pluripotent stem cell(iPSC)-derived neurons from patients with *GBA-*related PD and GD showed widespread ALP impairment [30-32].

A key regulator of the ALP is transcription factor EB (TFEB). In homeostatic conditions TFEB is retained in the cytosol by phosphorylation, primarily by the inhibitory action of mTORC1 [33, 34]. The presence of stress stimuli - such as starvation [35], ER-stress [35] and oxidative stress [36] - result in mTORC1 inhibition and dephosphorylation of TFEB, inducing its translocation into the nucleus. Nuclear TFEB increases the expression of genes involved in the regulation of lysosomal, autophagic and retromer function, collectively called Coordinated Lysosomal Expression and Regulation (CLEAR) network, which include *GBA* and *TFEB* itself [27, 33, 37-39]. Results from a pioneering study by Decressac *et al*. implicated TFEB in synucleinopathies, as nuclear localization of TFEB was reduced in a rat model overexpressing aSyn and in postmortem midbrain brain tissue sections of PD donors [40]. Moreover, both overexpression and direct pharmacological activation of TFEB protect against accumulation of aSyn/tau and associated neurodegeneration in several murine and in vitro models [41-44], suggesting TFEB could represent a potential therapeutic target for PD and other proteinopathies, such as DLB and Alzheimer’s disease [43, 45-47].

Here, we aimed at investigating the relevance of TFEB subcellular distribution and its relation to the presence of aSyn cytopathology in early (iLBD) and late stages PD/DLB, both sporadic (sPD/DLB) and *GBA*-related (*GBA*-PD/DLB). To this end, we explored the subcellular localization of TFEB in dopaminergic neurons in the substantia nigra pars compacta (SNpc) of post-mortem human brains using high-resolution confocal and stimulated emission depletion microscopy (STED) microscopy, accounting for the presence or absence of intracellular pSer129 aSyn deposition. We included post-mortem midbrain tissue of iLBD patients (N=3), PD/DLB patients with *GBA* variants (n=10), sporadic PD/DLB patents (N=9) and control subjects (N=7). iLBD cases were included as prodromal stages of PD/DLB [48].

Our findings showed reduced nuclear localization of TFEB in sPD/DLB in SNpc dopaminergic neurons, supporting previous findings [40], particularly in neurons with Ser129 aSyn+ cytopathology. Additionally, we observed an unanticipated clustering of TFEB which localized at the Golgi apparatus (GA), which were strongly increased in the sPD/DLB group compared to controls. The effect was more prominent in patients carrying *GBA* risk variants. Interestingly, TFEB clustering was increased in iLBD cases compared to controls. Remarkably, the increase in TFEB clusters was observed both in cells without pSer129 aSyn cytopathology as well as in neurons with intracellular pSer129 aSyn. Moreover, the TFEB immunopositive clusters did not colocalize with intracellular aSyn aggregates. Semi-quantitative measurement of the TFEB clusters in aSyn-negative cells associated with reduced total GCase enzymatic activity and with disease progression, as defined by Braak LB stages. Our results support a role of impaired TFEB localization in the early stages of disease pathogenesis in sporadic and *GBA*-PD/DLB, and suggest its involvement prior Ser129p aSyn accumulation

## Materials and methods

### Post-mortem human midbrain tissue

Post-mortem human brain tissue was obtained from neuropathologically-verified donors with iLBD (3 individuals), clinically diagnosed and neuropathologically-confirmed advanced sporadic PD or DLB without *GBA* variants (9 individuals), PD or DLB patients with *GBA* risk variants PD (9 individuals) and 15 age-matched control subjects without *GBA* risk variants. These patients represent a subset of a cohort for which the genotyping and various biochemical measurements were previously published [14, 49]. The demographics and group characteristics of the selected patients are presented in Table 1 and Table S1. *GBA* severe variants were identified according to their association with Gaucher disease (GD) type II or III, corresponding to *Parlar et al.* using the *GBA-PD browser* [50, 51], when available (Table S1). One patient with multiple *GBA* variants (p.Asp140His, p.Glu326Lys, and p.Thr369Met) was classified as severe.

**Table 1:** Group characteristics of controls, iLBD, sPD/DLB and GBA-PD/DLB patients in the present IHC study.

|  | **Controls**<br>**(N=7)** | **iLBD**<br>**(N=3)** | **sPD/DLB**<br>**(N=9)** | **GBA-PD/DLB**<br>**(N=9)** |
| --- | --- | --- | --- | --- |
| **Age of death (yrs ± SD)** | 77 ± 5 | 92 ± 6 | 77 ± 4 | 77 ± 6 |
| **Sex (M/F)** | 1/6 | 0/3 | 7/2 | 5/4 |
| **Postmortem delay (min ± SD)** | 342 ± 96 | 336 ± 84 | 330 ± 60 | 330 ± 90 |
| **Braak Lewy Body stage** | 0-1 (6/1) | 3 (3) | 4-6 (2/3/4) | 4-6 (2/2/5) |
| **Braak stage for**<br>**Neurofibrillary Tangles** | 1-2 (2/5) | 2-3 (2/1) | 0-2 (3/4/2) | 1-3 (8/0/1) |
| **CERAD Amyloid Plaque**<br>**Score** | O-B (1/4/2) | O-A (2/1) | O-B (4/2/3) | O-C (4/2/2/1) |
| **Thal phase** | 0-2 (1/2/3) | 0-1 (2/1) | 0-3 (4/0/1/2) | 0-4 (4/1/2/1/1) |
| **Total GCase enzymatic**<br>**activity in SN**<br>**(pmol/min/mg)[49]** | 1032 ± 142 | - | 828 ± 116 | 553 ± 209 |

Following all ethical and legal guidelines, informed consent for brain autopsy, the use of brain tissue, and clinical information for scientific research was given by either the donors or their family. Brains were dissected in compliance with standard operating protocols of the Netherlands Brain Bank and Brain Net Europe, and neuropathology was assessed by an experienced neuropathologist, according to the guidelines of Brain Net Europe [52, 53]. All donors were selected based on limited concomitant AD pathology (Braak neurofibrillary tangle stage ≤ 3 and CERAD ≤ B) and without micro-infarcts. For the groups of patients with iLBD, we selected cases with Braak LB stages 2 or 3, while the PD/DLB diagnostic group was selected with a Braak LB stage ≥ 4 [54]. Additionally, control subjects where selected based on the absence of significant aSyn pathology (Braak LB score ≤ 1), no record of neurological disorders and lack of any *GBA* risk variant. Postmortem delay (PMD) for all donors was less than 10 hours. Formalin-fixed paraffin-embedded (FFPE) tissue blocks of the midbrain containing the SN from all included donors were cut into 10μm thick sections, which were used for immunohistochemistry and multiple labelling experiments. Frozen SN and medial frontal gyrus (MFG) tissue was used for the mRNA quantification and GCase enzymatic activity quantification. For this, the tissue was pulverized frozen in pre-cooled stainless-steel grinding jars using a cryogenic grinder for 2 minutes or until full pulverization (30Hz, Mixer Mill MM400, Retsch, Haan, Germany).

### Immunohistochemical staining procedure

Double and triple labeling experiments were performed on 10μm thick FFPE sections. After deparaffinization, antigen retrieval was performed using 10mM citrate buffer (pH 6) in a steamer at 96°C for 30 minutes. In double labelling protocols, sections were subsequently blocked in a blocking buffer (BS) containing 2%normal donkey serum and 0.1% Triton-X in TBS (pH 7.6) for one hour, after which primary antibodies were incubated overnight at 4°C. Afterwards, the sections were washed and incubated in appropriate secondary antibodies diluted in BS for 2 hours at room temperature. In case of multiple labelling experiments, after incubation with secondary antibodies sections were blocked for 1 hour in 5% normal rabbit serum and 5% normal mouse serum in TBS, and incubated for 2 hours at room temperature in BS containing directly labelled antibodies [4]. Direct labelling of antibodies was done using a Zenon™ Alexa FluorTM 594 Rabbit IgG Labelling Kit (# Z25307, ThermoFisher Scientific). DAPI was added to the BS in a concentration 1 µg/ml in the last incubation step. Sections were mounted in Mowiol mounting solution using glass cover slips (Art. No.: 630-2746; Glaswarenfabrik Karl Hecht, Sondheim, Germany). Negative control stainings lacking primary antibodies were performed to control for background/autofluorescence levels and a-specific staining. Single labelling for each antibody included in multiple-labelling experiments were carefully examined to control whether immunoreactivity patterns were caused by possible cross-reactivity between antibodies. The specificity of the TFEB antibodies used in this study was confirmed by WB (Fig. S1.B) [43, 55]. The antibodies utilized for the experiments of this study are summarized in Table S2.

### Confocal and stimulated emission depletion microscopy

Confocal laser scanning microscopy (CSLM) and stimulated emission depletion (STED) microscopy were performed using a Leica TCS SP8 STED 3X microscope (Leica Microsystems). All images were acquired using a HC PL APO CS2 100x 1.4 NA oil objective lens, with the resolution set to a pixel size of 20 nm x 20 nm. Gated hybrid detectors were used in counting mode. Sections were sequentially scanned for each fluorophore, by irradiation with a pulsed white light laser at different wavelengths. A pulsed STED laser line at a wavelength of 775 nm was used to deplete the Alexa594 fluorophore, while a continuous wave (CW) STED laser with wavelength of 592 nm was used to deplete the Alexa 488 fluorophore, respectively. After scanning, deconvolution was performed using CMLE (for confocal images) or GMLE (for STED images) algorithms in Huygens Professional software (Scientific Volume Imaging, Huygens, the Netherlands). All images were adjusted for brightness/contrast in the same way using an ImageJ (National Institute of Health, USA) script before image analysis.

### Single-cell scanning and semi-quantitative scoring

Images of neuromelanin-containing cells in the SNpc were collected using a 100x 1.4 NA oil objective lens. The neuromelanin-containing portion of the cytoplasm was defined for each neuron using brightfield scans. In addition, selection of cells for scanning was based on the visibility of a nucleus and a defined cytoplasmic portion not filled with neuromelanin granules in the same z-plane. The presence of somatic pSer129 aSyn immunoreactivity was used to distinguish between cells with and without cytopathology according to previously defined subcellular phenotypes of pSer129 aSyn pathology. TFEB reactivity patterns were not used as a criterion for selection. For each cell, a z-stack of 1.05µm was scanned based on the z-position of the nucleus and when possible nucleolus) using a z-step size of 0.15µm. A total of 441 neuromelanin-containing dopaminergic cells was scanned, including 100 neurons of age-matched control subjects, 105 of iLBD, 109 of sPD/DLB and 129 of *GBA*-PD/DLB patients (Table S3). The number of imaged cells that matched our defined inclusion criteria ranged from 7 to 39 per individual and was restricted by extensive loss of nigral neurons in PD/DLB patients. A distinction was made between neurons with and neurons without subcellular immunoreactivity for pSer129 aSyn. This was done according to the previously described detailed subcellular phenotypes of pSer129 aSyn – including different cytoplasmic inclusions as well as cytoplasmic network. As these features were not observed in controls in a previous study by us [4] or in the present study, they were therefore considered as highly specific for aSyn cytopathology.

Raw confocal images, which were scanned using the same settings, were processed in the same way in ImageJ. For every scanned cell, the images from brightfield, TFEB and DAPI channels were extracted for semi-quantitative analysis, without information about pSer129 aSyn. Subsequently, all images were blinded for diagnosis by randomized coding and nuclear TFEB reactivity was semi-quantitatively assessed by two independent raters. Based on pilot measurements in a subset of cells, the following scores were defined: 1 – a pronounced lower density of punctate TFEB in the nucleus versus cytoplasm; 2 – comparable densities of punctate TFEB in the nucleus versus cytoplasm; 3 – pronounced nucleolar labeling. The inter-rater reliability of the nuclear TFEB scoring, estimated by the Cronbach’s alpha coefficient, was 0.88. An unexpected feature of TFEB immunoreactivity patterns was the presence of cytoplasmic clusters, seemingly independent of cytoplasmic TFEB punctae (discussed in results section). This feature was semi-quantitatively scored in the same images by two independent assessors as follows: 0 – no clusters, 1 – low (1-2-clusters), 2 – intermediate (< 10 clusters), 3 – severe (> 10 clusters). The inter-rater reliability of the TFEB cluster scoring, as estimated by the Cronbach’s alpha coefficient, was 0.91.

### RNA extraction and mRNA quantification

Messenger RNA expression levels of selected CLEAR genes were analyzed using quantitative polymerase chain reaction (qPCR). For this purpose, total RNA was extracted from pulverized SN as well as MFG in a subset of cases, depending on tissue availability (see Table S1). A Trizol/chloroform protocol for RNA extraction was used, as previously published [56]. To check for RNA quality, the purity of the extracted RNA was estimated based on the ratio of absorbance at 260/280 nm λ using a Nanodrop spectrophotometer (ThermoFischer Inc., Waltham, MA, USA). RNA integrity was calculated based on the 28S to 18S rRNA ratio using the RNA 6000 Nano Kit and the Agilent 2100 Bioanalyzer (Agilent Technologies Inc., Santa Clara, CA, USA) and was expressed as RNA integrity number (RIN). Only samples with RIN values ≥ 5.0 and PMD < 10 hours were included in the analysis. SN samples had RIN values between 5.0 and 7.9 (mean=6.0, variance=0.48) and MFG samples between 5.6 and 8.0 (mean=6.6, variance=0.46). From the extracted total RNA samples, cDNA was synthetized using the High-Capacity cDNA Reverse Transcription Kit (Applied Biosystems, cat. n. 4368814, ThermoFischer Inc., Waltham, MA, USA). Negative controls lacking the RT enzyme were included in the procedure to control for genomic DNA contamination during qPCR.

Intron-spanning primers were designed for the selected genes encoding for galactosylceramidase (*GALC*), hexosaminidase subunit alpha (*HEXA*), ß-glucocerebrosidase (*GBA*), microtubule associated protein 1 light chain 3 (*MAP1LC3A*), sequestosome 1 (*SQSTM1*), UDP-glucose ceramide glucosyltransferase (*UGCG*), VPS35 retromer complex component (*VPS35*) and for the reference genes pescadillo ribosomal biogenesis factor 1 (*PES1*), RNA polymerase II subunit A (*POLR2A*), ornithine decarboxylase antizyme 1 (*OAZ1*) and RNA polymerase II subunit F (POLR2F). Suitable TaqMan probes were selected from the Universal Probe Library (Roche Applied Science, Indianapolis, IN, USA). All primer used were purchased from Kaneka Eurogentec (Seraing, Belgium) and are listed in Table S4. Each cDNA sample was analyzed in triplicate (mean CV=0.4%) and standard curves and efficiency values were calculated using the StepOne plus software (Thermo Fisher). The software geNorm (Genorm) was used to select the reference genes with the most stable expression among all cases. Results were quantified as relative expression normalized on the geometric mean [57, 58] of most suitable reference genes (PES1, POLR2A and POLR2F).

### GCase enzymatic activity quantification

Selected cases included in this study are part of a larger cohort for which bulk-tissue glucocerebrosidase activity was measured in [49] and the values relative to subset of cases included in the present study were extracted. Briefly, pulverized MFG and SN tissue from a subset of cases were lysed in 50 mM sodium/phosphate (Na/P), 150 mM NaCl buffer, pH 7.0, homogenized, sonicated and centrifuged as previously described [49]. Protein concentration was determined using a Bradford assay [59] and aliquots containing equal amounts of protein were incubated at 37°C with 3mM of the florigenic substrate 4-methylumbelliferyl-β-D-glucopyranoside (MUB-Glc) in equal amounts of MCIlvaine reaction buffer (0,1m citric acid, 0,2 m Na_2_HPO_4_), pH4.5 with 0.2% taurodeoxicolate. After 60 minutes, the reaction was stopped with ice-cold 0.2 M glycine-NaOH buffer, pH 10.4 and the fluorescence of MUB released by enzymatic cleavage was measured with a FLUOstar Optima microplate reader (BMG Labtech, Ortenberg, Germany [λex/λem=360nm/450nm]) [60]. Each sample was analyzed in triplicate alongside a negative control condition lacking the protein sample and an unbounded MUB standard curve. The enzymatic activity was calculated based on the MUB standard curve and expressed as amount of total protein necessary to hydrolase 1 nmol of MUB-Glc in 1 minute at 37°C (pmol/min/mg). The enzymatic activities measured in the cases selected for this study showed a mean CV of 4.1% (variance=3.3) and of 4.9% (variance=4.3) for SN and MFG, respectively.

### Statistics and computing

In order to analyze the TFEB semi-quantitative scores, we applied Generalized Estimating Equations (GEE) models with an ordinal probit response for our statistical analysis to assure that our results could not be attributed to measurements in a single individual or to differences in the number of pathological cells per individual. For each test, Wald Chi-Square and B statistics (together with 95% confidence interval) are presented in the results section. For correlation analyses, either Pearson’s or Spearman’s tests were performed as required and as indicated in the results section. When comparing the normally-distributed continuous data from the GCase enzymatic activity assay, one-way ANOVA with group as fixed factor and age as covariate was used to test for differences between the groups’ means, coupled with Games-Howell post-hoc test to identify specific differences between the groups. For the analysis of the mRNA quantification data, General Linear Models (GLM) were used with gene and disease group as fixed factors and age as covariate. All statistical analyses were performed using the software IBM SPSS statistics (version 26.0, IBM, Armonk, NY, USA). In all tests, significance threshold was set a p ≤ 0.05. Graphs were generated using the software GraphPad Prism (version 8.2.1, GraphPad Software, San Diego, CA, USA). The graphical abstract (Fig. 7) was created using Biorender (BioRender.com, 2022).

## Results

### Description of observed TFEB immunoreactivity patterns in SNpc neurons

When analyzing TFEB immunostaining patterns using the selected antibodies (Fig. 1.1,S1,S2,S3) in iLBD, sporadic and *GBA*-related PD/DLB, we observed heterogeneous profiles and distribution patterns in neuromelanin-containing dopaminergic neurons in the SNpc in all cases. Such heterogeneity could be observed among the neurons of each subject (Fig. S2).

**Figure 1:**
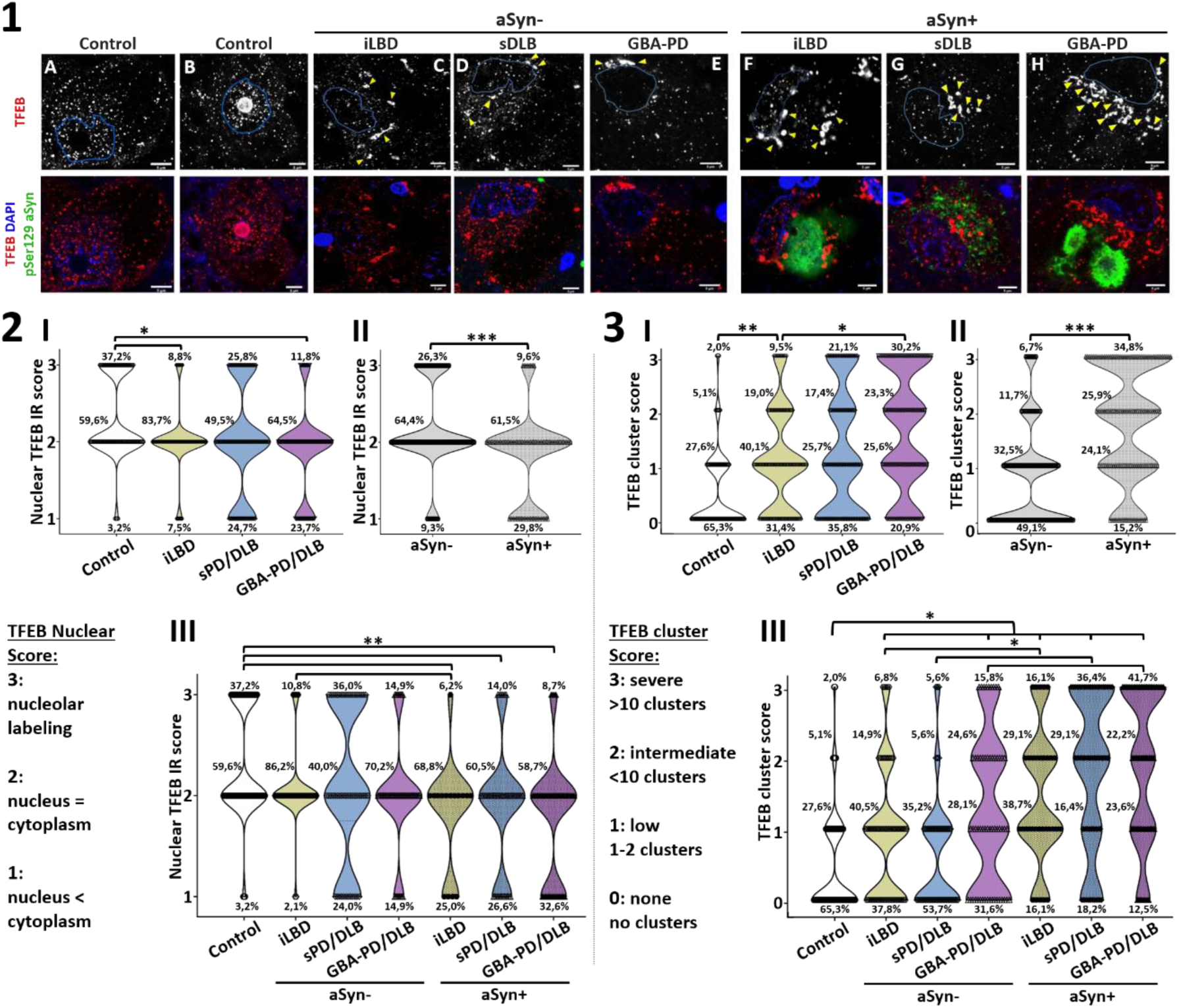
Reduced nuclear localization and increased perinuclear clustering of TFEB in iLBD, sPD/DLB and GBA-PD/DLB compared to controls in dopaminergic SN neurons with and without pSer129 aSyn cytopathology. **1)** Representative raw confocal images of TFEB localization patterns in dopaminergic SNpc neurons in iLBD, PD/DLB patients and controls. **A-B:** Neurons from control cases. **B-E:** Neurons without pSer129 aSyn cytopathology; **F-H:** Neurons with pSer129 aSyn cytopathology. Features immunoreactive for TFEB included nuclear and cytoplasmic punctae in different proportions, nucleolar staining patterns (*A*), and perinuclear clusters (*B-H*). The outline of the DAPI signal is projected in blue. Yellow arrow heads indicate TFEB immunopositive clusters. All scale bars represent 5µm. **2)** Nuclear TFEB is reduced in the disease groups compared to controls. **I:** Frequency distributions of nuclear TFEB immunoreactivity (IR) semi-quantitative scores (as indicated). **II:** Frequency distribution of nuclear TFEB IR scores in cells with and without subcellular pSer129 aSyn pathology across all diagnostic groups. **III:** Frequency distributions of nuclear TFEB IR scores in cells with and without pSer129 aSyn pathology per group. Higher nuclear TFEB IR scores were more frequently found in controls compared to the iLBD, sPD/DLB and GBA-PD/DLB groups. **3)** TFEB clusters are increased in the disease groups both in presence and absence of pSer129 aSyn cytopathology. **I:** Frequency distributions for semi-quantitatively scored TFEB clusters (as indicated). **II:** Frequency distribution of TFEB cluster scores in cells with and without subcellular pSer129 aSyn pathology across all diagnostic groups. **III:** Frequency distributions of TFEB cluster scores in cells with and without pSer129 aSyn pathology per disease group. An increase in the TFEB cluster score was observed in iLBD, sPD/DLB and GBAPD/DLB compared to controls, both in cells with and without pSer129 aSyn pathology. Within each disease group, TFEB clusters were observed more frequently in cells with pSer129 aSyn cytopathology compared to cells without pSer129 aSyn cytopathology. aSyn+: neurons with pSer129 aSyn pathology; aSyn-: neurons without pSer129 aSyn pathology. *p < 0.05; ** p < 0.005; *** p < 0,001.

### -Nuclear and cytoplasmic TFEB punctae

Most prominently, immunopatterns were characterized by many TFEB-positive punctae, which were distributed throughout the cytoplasm as well as in the nucleus (Fig. 1.1, Fig. S2). In a subset of these neurons, the nucleolar portion of the nuclear compartment – outlined based on DAPI signal – was strongly labeled by TFEB (Fig. 1.1-B). Overall, the scanned neurons showed heterogeneity in the distribution of TFEB immunopositive punctae in the nucleus versus the cytoplasm ranging from neurons displaying high density of TFEB punctae in nucleus compared to cytoplasm (Fig. 1.1-A,B) to neurons with barely any nuclear TFEB punctae (Fig. 1.1-E, 1.1-H).

### -Somatic TFEB clusters

In addition to immunoreactive punctae (typically around 0.3µm in the raw CSLM signal), a subset of neurons (particularly in disease groups) displayed larger TFEB immunoreactive clusters in the perinuclear cytoplasm (Fig. S2-S3, Fig. 1.1-C,D,F,G,H: yellow arrowheads). These clusters appeared more frequently in neurons from diseased patients compared to controls and could be observed both in neurons without intracellular aSyn as well as in neurons with cytopathology. The frequency and number of the observable cytoplasmic clusters appeared variable among neurons and seemed to be increased in the presence of subcellular pSer129 aSyn immunoreactivity compared to neurons without apparent aSyn cytopathology (Fig. 1.1-F,G,H). These clusters were observed using different TFEB antibodies (Fig. S3).

To verify that these profiles were not the result of non-specific antibody binding in fixed tissue specimens and can be also observed in vitro, we differentiated hESC-derived neurons from a healthy control case (wt) and an isogenically-derived hESC line lacking *GBA* (*GBA* KO) to model for the reduced GCase activity observed in *GBA*-PD/PDD. After staining for TFEB and cytoskeletal protein MAP2, a CLSM analysis revealed the presence of a subset neurons (both WT and *GBA*-KO) displaying perinuclear immunopositive TFEB clusters, similar to what observed in vivo (Fig. S1-A). The frequency of cells with TFEB-positive clusters did not differ between the two lines. Interestingly, the neurons displaying TFEB clusters often displayed signs of cellular stress, such as altered cellular morphology, reduced nuclear shape and DNA condensation, suggesting this phenotype might be associated with cellular stress.

### Nuclear TFEB immunoreactivity is reduced in iLBD, sPD/DLB and GBA-PD/DLB

In order to compare nuclear TFEB between controls and the different disease groups, the nuclear TFEB immunoreactivity was semi-quantitatively scored in a blind analysis (see *Materials and methods*). The analysis showed an increased number of neurons with nuclear TFEB in controls compared to the diseased groups (p=0.025; Wald Chi-Square: 4.995; B=0.754; 95% confidence interval (CI) [0.093:-1.415]). In particular iLBD donors (p=0.043, Wald Chi-Square: 4.093; B=-0.665; 95% confidence interval (CI) [-1.308:-0.021]) and *GBA*-PD/DLB patients (p=0.009; Wald Chi-Square: 6.750; B=-0.927; 95% confidence interval (CI) [-1.627:-0.228]) showed reduced nuclear TFEB immunoreactivity compared to controls, while this effect was less pronounced in advanced sPD/DLB patients (p=0.113; Wald Chi-Square: 2.513; B=-0.669; 95%CI [- 1.496: 0.158]; Fig. 1.2-I). These results are in support of previous indications that nuclear TFEB immunoreactivity is reduced in dopaminergic neurons in the SNpc of patients with PD compared to controls [40].

Interestingly, the results of our analyses convincingly showed that nuclear localization of TFEB was reduced in neurons affected by pSer129 aSyn cytopathology compared to neurons without intracellular pSer129 aSyn immunoreactivity (Fig. 1.2-II). When comparing neurons with and without pathology scanned in the different diagnostic groups, we observed less nuclear immunostaining in cells with pSer129 aSyn pathology than without (p=0.0005; Wald Chi-Square: 11.959; B=-0.720; 95%CI [- 1.128:-0.312]). This effect was still significant when excluding controls from the analysis (p=0.0013; Wald Chi-Square: 10.308; B=-0.521; 95%CI [-0.839:-0.203]). While nuclear TFEB immunoreactivity in neurons with pSer129 aSyn cytopathology was reduced compared to controls both in patients with either iLBD, sPD/DLB and *GBA*-PD/DLB (Fig. 1.2-III), we observed a trend for lower nuclear immunoreactivity scores also in neurons without apparent aSyn pathology in the iLBD group (p=0.104; Wald Chi-Square: 2.646; B=-0.543; 95%CI [-1.197:0.111]) and in the *GBA*-PD/DLB group (p=0.071; Wald Chi-Square: 3.253; B=-0.705; 95%CI [-1.470:0.061] compared to cells in control. This trend was not observed in sPD/DLB patients (p=0.365; Wald Chi-Square: 0.821; B=-0.465; 95%CI [-1.472:0.541]).

### Cytoplasmic clustering of TFEB is increased in iLBD, sPD/DLB and GBA-PD/DLB

The semi-quantitative scores of TFEB clusters in the cytoplasm of neurons showed pronounced differences between diagnostic groups. The proportion of neuron displaying TFEB clusters was significantly higher in diseased patients compared to controls (p=0.0002; Wald Chi-Square: 14.019; B=1.034; 95% CI [0.493:1.575]). This effect was significant in iLBD (p=0.004; Wald Chi-Square: 8.218; B=0.798; 95% CI [0.252:1.343]), in sPD/DLB (p=0.004; Wald Chi-Square: 8.227; B=0.938; 95% CI [0.297:1.580]) as well as in *GBA*-related PD/DLB patients (p=0.0003; Wald Chi-Square: 17.447; B=1.336; 95%CI [0.709:1.962]) compared to controls, with the latter being the group with the strongest effect (Fig. 1.3-I). Strikingly, TFEB clusters were increased both in cells with and without aSyn cytopathology (Fig. 1 and Fig. 2). The effect was especially pronounced for aSyn-negative cells of the *GBA*-PD/DLB group, where a 8-fold increase in the percentage of cells with severe TFEB clusters (score 3) was observed compared to controls.

**Figure 2:**
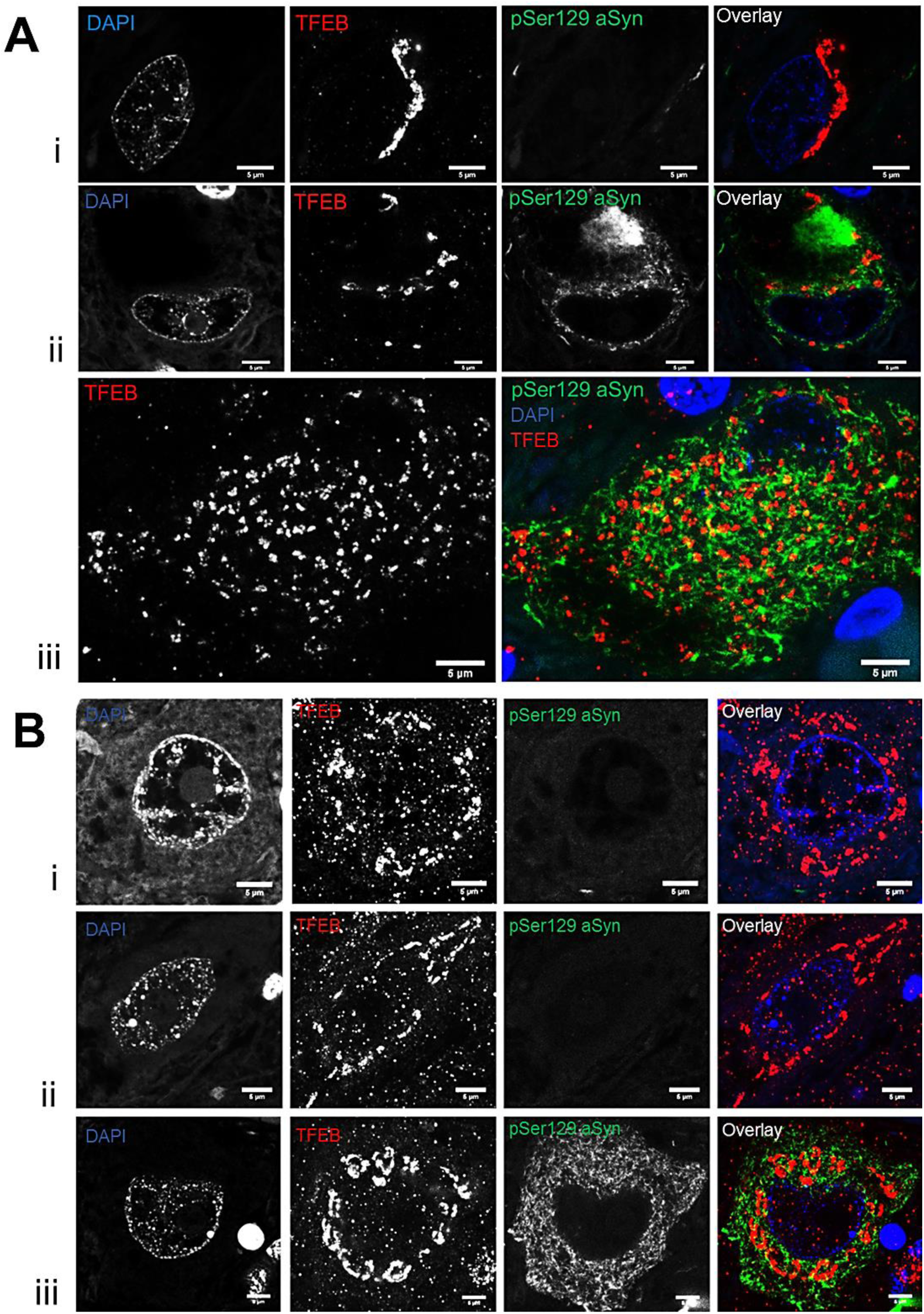
Representative TFEB immunoreactivity patterns in dopaminergic neurons in the SNpc of patients carrying severe *GBA* variants. Representative confocal images are shown for donor ID26 with an L444P variant (**A**) and for donor ID24, which carries multiple *GBA* variants (**B**). TFEB immunoreactivity in neurons of PD patients with severe GBA variants was characterized by dense clusters in the cytosol, with limited immunopositive punctae in the cytoplasm and, particularly, in the nucleus. These patterns were not only observed in neurons with pSer129 aSyn pathology (**Aii**, **Aiii**; **Biii**), but also in neurons without apparent aSyn pathology (**Ai; Bi, Bii**). All scale bars represent 5µm.

The amount of clusters was particularly increased with the presence of pSer129 aSyn cytopathology, as cells that contained pSer129 aSyn pathology showed more TFEB clusters compared to cells without pathology (p= 2.38E-12; Wald Chi-Square: 49.139; B=1.092; 95%CI [0.787:1.558], Fig. 1.3-II). The increase remained significant when excluding controls from the analysis (p=8.77E-11; Wald Chi-Square: 42.078; B=0.892; 95%CI [0.623:1.162]).

However, our analysis showed an increase in the TFEB clustering also in cells without pSer129 aSyn in diseased groups compared to controls. Neurons negative for pSer129 aSyn showed a significantly more TFEB clustering in iLBD (p=0.016; Wald Chi-Square: 5.756; B=0.649; 95%CI [0.119:1.180]) and *GBA*-PD/DLB (p=0.018; Wald Chi-Square: 5.572; B=0.991; 95%CI [0.168:1.814) patients compared to controls (Fig. 1.3-III), while no statistically significant difference was observed in cells of sPD/DLB patients without cytoplasmic pSer129 aSyn immunoreactivity (p=0.421; Wald Chi-Square: 0.647; B=0.295; 95%CI [-0.424:1.013]).

### Cytoplasmic clustering of TFEB in patients with pathogenic GBA variants

Our semi-quantitative analyses alterations in TFEB distribution were most extreme in the group of *GBA*-PD/DLB patients compared to controls. To further explore the impact of *GBA* mutations on TFEB, we selected two donors of our cohort with severe *GBA* mutations to assess TFEB immunoreactivity patterns in more detail. TFEB immunopatterns in the patients with pathogenic *GBA* variants and resulting GCase deficiency revealed severe TFEB clustering in neuromelanin-containing dopaminergic neurons (Fig. 2). The effect was observed both in cells with (Fig. 2.A-i,B-i) and without pSer129 aSyn cytopathology (Fig. 2.A-ii, 2.B-ii). In these cells, TFEB immunoreactivity was concentrated in large cytosolic clusters showing limited additional immunopositive punctae in the cytoplasm and in the nucleus and these effects appeared more extreme compared to iLBD and sPD/DLB patients, as well as compared to patients with milder *GBA* variants.

### TFEB clusters at the Golgi apparatus

TFEB immunopositive clusters were localized mainly at the perinuclear portion of the cytoplasm. We used STED microscopy to visualize the clusters in more detail and observed that the arrangement of TFEB within the clusters resembled a cisternal morphology. This led us to the hypothesis that TFEB might cluster at the Golgi apparatus. This hypothesis was explored using a multiple labeling protocol with antibodies against pSer129 aSyn, TFEB, TGN46 – a marker for the trans-Golgi network (TGN) - and DAPI (Fig. 3) combined with super-resolution STED microscopy. Our results demonstrated the localization of the described TFEB clusters to the Golgi (Fig. 3). Where in most neuromelanin-containing dopaminergic neurons, smaller TFEB clusters occupied a small part of the Golgi (Fig. 3.A,C), in other cells extensive TFEB labelling was observed throughout the entire structure (Fig. 3.B). In neurons without clusters, cytoplasmic TFEB-immunopositive punctae revealed only limited colocalization with TGN46 (Fig. 3.D). Colocalization of TFEB clusters at the Golgi was also observed using the cis-Golgi marker GOLGA2 (Fig. S4). Together, our STED observations suggests that TFEB accumulates at the Golgi under pathological conditions in sporadic and *GBA*-PD/DLB.

**Figure 3:**
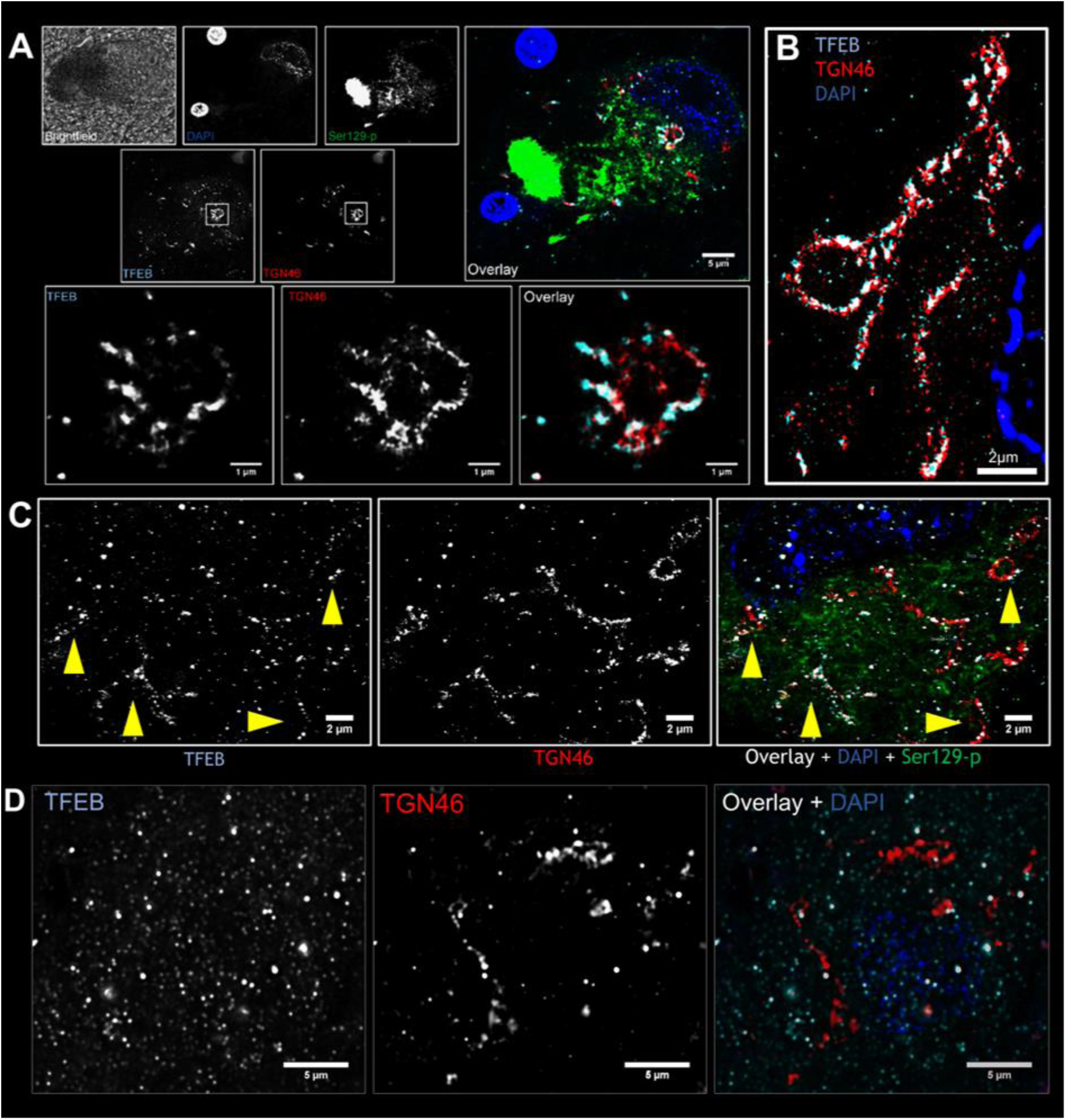
Cytoplasmic clusters immunoreactive for TFEB localize at the Golgi in dopaminergic SNpc neurons. Deconvolved confocal (**A** upper panels and **D**) and STED (**A** lower panels, **B**, **C**) images. **A:** Overview image (upper panels) and detailed magnification (lower panels) of the co-localization of TFEB clusters at the Golgi (visualized by TGN46), taken in sPD patient ID17, in a dopaminergic SNpc neuron with subcellular pSer129 aSyn pathology. **B:** Image taken in a dopaminergic SNpc neuron with subcellular pSer129 aSyn pathology in GBA-PD patient ID 24, demonstrating extensive perinuclear TFEB immunoreactivity localizing in the Golgi and displaying a detailed cisternal morphology (the pSer129 aSyn channel is not shown here). **C:** Image taken in a DLB patient with a Glu326Lys variant (ID37) showing TFEB immunopositive punctae in the cytoplasm and partial co-localization to the Golgi (indicated by yellow arrow heads). **D:** Image taken in a dopaminergic neuron of a control (ID9) without aSyn cytopathology, in which cytoplasmic TFEB immunopositive punctae show very limited co-localization with TGN structures. Scale bars: **A**, upper panels = 5µm; lower panels = 1µm; **B:** 2µm; **C:** 2µm; **D:** 5µm

### TFEB and TFEB clusters do not co-localize with aSyn

Altered subcellular TFEB localization was observed both within neurons with intracellular deposits of aSyn, being a LB, an amorphous aggregate or a diffuse intracellular staining, as well as within neurons without appreciable intracellular aSyn accumulation (Fig. 4.1). Nonetheless, aSyn immunopositive neurons showed a higher number of clusters overall compared to aSyn-negative cells (Fig. 1). Previous reports have described potential protein-protein interaction between TFEB and aSyn [61, 62]. To investigate whether aSyn could be causing TFEB mislocalization by direct interaction, we investigated weather co-localization of the two proteins could be observed in SNpc neurons with intracellular aSyn accumulation in sPD/DLB and *GBA*-PD/DLB cases. First, we stained for TFEB and for aSyn with an antibody directed towards the N-term portion of the protein (aa 1-60, N-19) and expected to recognize a substantial pool of non-truncated and truncated aSyn proteoforms (Fig. 4.1) [63]. CLSM and STED microscopy showed no colocalization between TFEB clusters and aSyn (Fig. 4.1-A). Similarly, analysis of TFEB and pS129 aSyn immunostaining revealed only sporadic colocalization (Fig. 4.1-B, arrowheads). Together, these results suggest that TFEB clustering is not primarily due to a direct interaction with aSyn aggregates.

**Figure 4:**
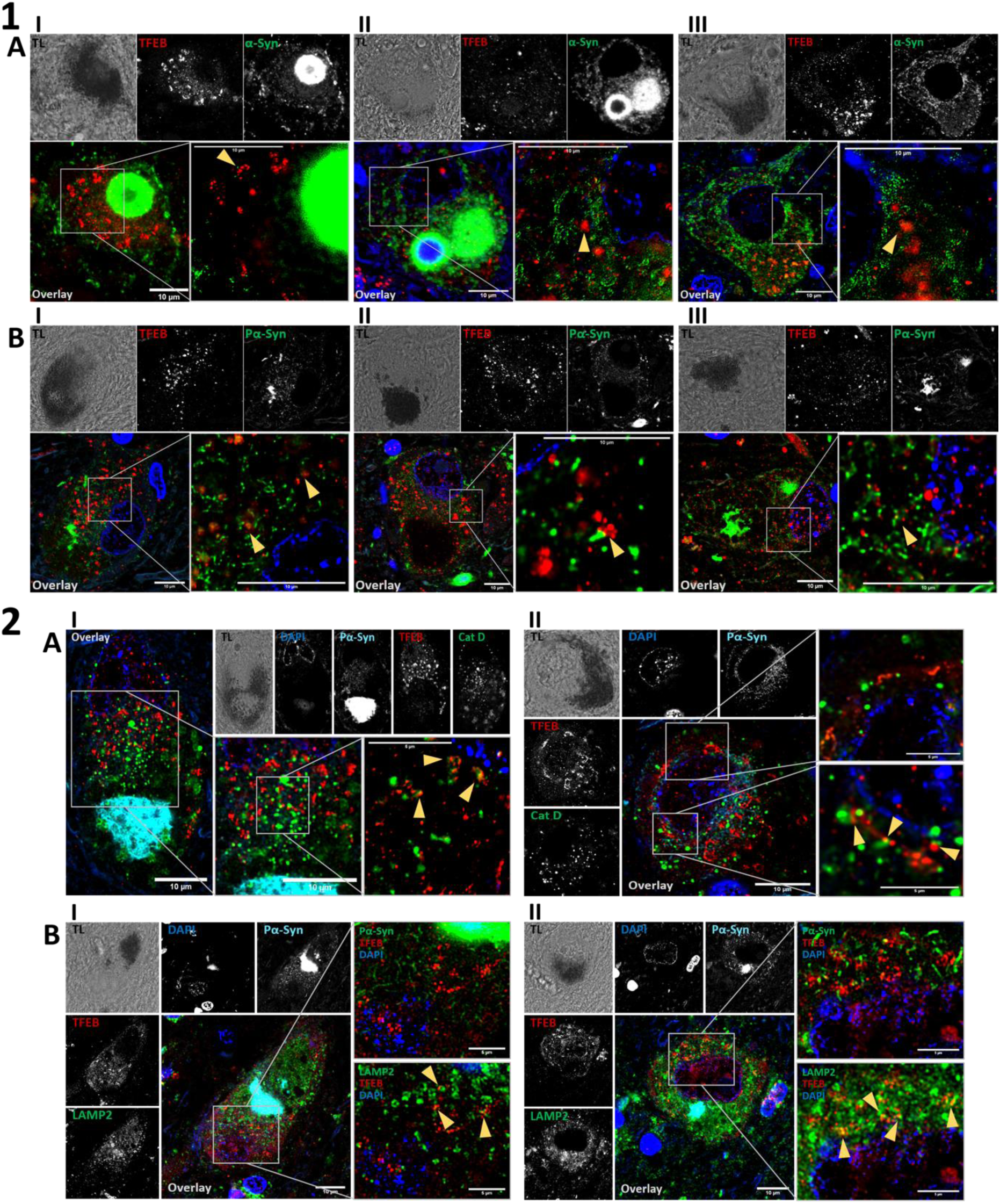
Cytoplasmic clusters immunoreactive for TFEB do not co-localize with alpha-synuclein and interact with lysosomes. Representative transmitted light (TL, upper-left panels), deconvolved CLSM (central and right upper panels) and STED (lower-right panels) images of dopaminergic SNpc neuron from donor ID27 (sPD, 1*-AI,BI*), ID37 (GBA-DLB, *1-AII*,*BII*) and ID16 (GBA-PD, *1-AIII,BIII*), ID43 (GBA-DLB, *2-AI*), ID39 (sDLB, *2-AII*)), ID22 (sPD, *2-BII*) and ID24 (GBA-PD, *2-BII*). **1)** Colocalization analysis of TFEB clusters with aSyn. **A**: Staining for TFEB (red) and alpha-synuclein (N-19, aSyn aa 1-60) (α-Syn, green) in neurons with intracellular LB (*I*), intracellular LB with amorphous aggregate (*II*) and diffuse intracellular staining (*III*) showing no co-localization with TFEB clusters. **B:** Staining for TFEB (red) and Ser129-phosphorylated alpha-synuclein (PS129 α-Syn, green) in neurons with intracellular small amorphous aggregate (*I, III*) and intracellular diffuse staining (*II*) showing limited co-localization with TFEB clusters. **2)** Colocalization analysis of TFEB clusters with lysosomes. **A:** Staining for TFEB (red), Ser129-phosphorylated aSyn (Pα-Syn, cyan) and lysosomal protein Cathepsin D (Cat D, green) in neurons with intracellular amorphous aggregate (*I*) and diffuse intracellular staining (*II*). **B:** Staining for TFEB (red), Ser129-phosphorylated aSyn (Pα-Syn, cyan/green as indicated in each panel) and lysosomal protein LAMP2 (LAMP2, green) in neurons with intracellular amorphous aggregate (*I* and *II*). The stainings show interaction between TFEB and TFEB clusters with the lysosomal pool. Scale bars = 5/10 uM µm as indicated in each panel.

### TFEB clusters retain the ability of interacting with the lysosomal pool

The main regulation of TFEB happens at the lysosomal level, as functional TFEB transiently associates with the cytoplasmic side of the lysosomal membrane where it is phosphorylated. To investigate whether the mislocalized TFEB observed in the perinuclear clusters retains the ability of interacting with lysosomes, we studied the co-localization of TFEB clusters with lysosomal markers by CLSM and STED microscopy (Fig. 4.2). Staining for the lysosomal luminal protein Cathepsin D (CTSD) (Fig. 4.2-A) and for the lysosomal membrane protein LAMP2 (Fig. 4.2-B) showed the presence of lysosome-TFEB contact points. Lysosomes sporadically interacting and colocalizing with the TFEB clusters, as well as with TFEB puncta could observed (Fig. 4.2: arrowheads). This suggest that the TFEB protein clustering at the Golgi apparatus might retain its functional activity and thus still interact with the lysosomal pool.

### Total GCase enzymatic activity and expression of selected CLEAR genes are reduced in sPD/DLB and GBA-PD/DLB compared to controls

We further investigated how cytoplasmic retention and clustering of TFEB in the diseased groups associated with the enzymatic activity of the lysosomal hydrolase GCase. We previously reported a decreased GCase activity in the SN of sPD/DLB and – more pronounced – of *GBA*-PD/DLB patients compared to controls in the same cohort [49]. In the subset of selected PD/DLB cases for this study, GCase activity was significantly decreased in patients compared to controls (mean difference=-253.4; p=0.0003; 95% CI [-376.0:-130.8]), and further reduced in *GBA*-PD/DLB compared to sPD/DLB (mean difference=-275.2; p=0.012; 95% CI [-486.4:-64.0]) (Fig. 5.A).

**Figure 5:**
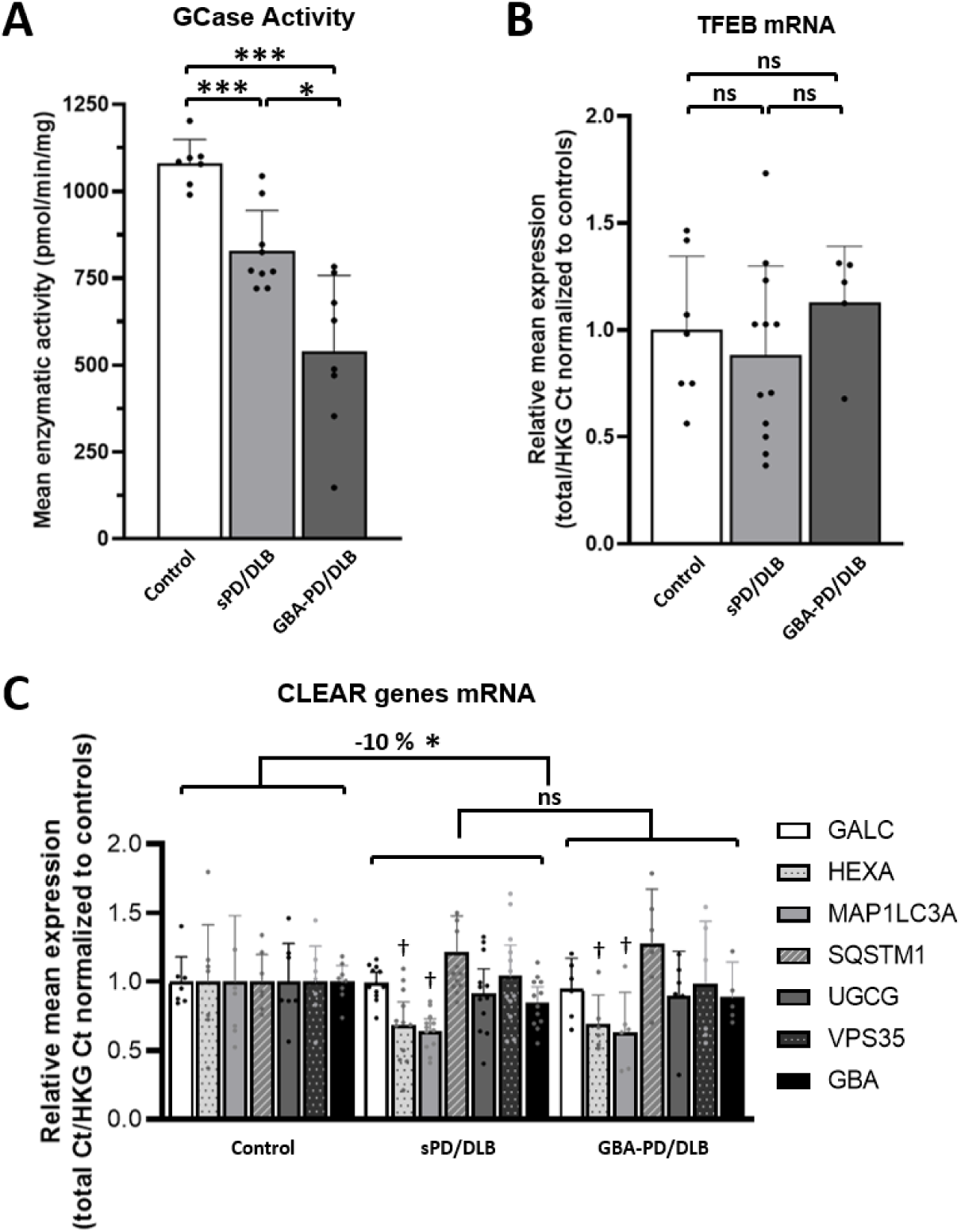
Impaired TFEB localization is associated with a reduction of bulk tissue total GCase activity and expression of selected CLEAR genes in the SN of sPD-DLB and GBA-PD/DLB patients. **A:** Total GCase enzymatic activity quantification in bulk SN tissue from sPD/DLB, GBA-PD/DLB patients and controls as measured in [49] and expressed as pmol/min/mg of total protein. Median ± 95% CI; N≥7/group, n=3. **B, C:** mRNA quantification by qPCR in sPD/DLB, GBA-PD/DLB patients and controls calculated as total Ct normalized on HKGs and expressed as fold-change compared to the control group. **B:** Quantification of TFEB mRNA. Mean ± SD; N≥5/group, n=3. **C:** CLEAR genes mRNA quantification. A 10% overall reduction in mRNA expression in the selected genes is observed when comparing the diseased group (sPD/DLB + GBA-PD/DLB) to controls. Mean ± SD. N≥5/group, n=3. HKG: housekeeping genes; GALC: galactosylceramidase; HEXA: hexosaminidase subunit alpha; GBA: ß-glucocerebrosidase; MAP1LC3A: microtubule associated protein 1 light chain 3; SQSTM1: sequestosome 1; UGCG: UDP-glucose ceramide glucosyltransferase; VSP35: retromer complex component. *p < 0.05; ***p < 0.001; † p < 0.05 vs Control.

To study whether the deregulation of TFEB observed could be associated with alterations in its transcription levels, we quantified TFEB mRNA by qPCR in sporadic and *GBA*-related PD/DLB and controls. Results did not show a statistically significant difference in the expression of the TFEB gene in the disease groups compared to controls (Fig. 5.B). Similarly, we quantified the expression of selected genes belonging to the CLEAR gene network which are under the transcriptional regulation of TFEB [38]. We observed a modest (10%), statistically significant collective reduction in the expression of the selected CLEAR genes in the disease patients compared to control (P=0.048) (Fig. 5.C). This suggest a downregulation of the CLEAR gene network in the SN of patients with PD/DLB. Data obtained in MFG material showed a similar pattern (Fig. S5). No statistically significant difference was shown when testing for differences between individual diagnostic groups. Among the tested genes, MAP1LC3A showed reduction in mRNA expression in sPD/DLB (p=0.027; Wald Chi-Square: 4.899; B=0.366; 95%CI [0.042:0.691]) and *GBA*-PD/DLB (p=0.012; Wald Chi-Square: 6.384; B=0.357; 95%CI [0.080:0.635]) group compared to controls. For HEXA (p=0.017; Wald Chi-Square: 5.715; B=0.313; 95%CI [0.056:0.571]) and *GBA* (p=0.044; Wald Chi-Square: 4.065; B=0.140; 95%CI [0.004:0.275]) mRNA expression was found to be decreased only when comparing diseased patients grouped against controls (Fig. 5.C).

### TFEB cluster score correlates with bulk tissue total GCase activity and disease progression

As TFEB is a master regulator of GCase expression, we hypothesized that the observed cytoplasmic retention and clustering of TFEB could be associated with changes in total GCase enzymatic activity. To assess this, we conducted a Spearman’s correlation analysis between total GCase activity and mean TFEB cluster score value per case (Fig. 6.A). The analysis revealed a strong negative correlation between the two measurements (r=-0,523, p<0,01) in aSyn-negative cells, with cells from cases with lower total GCase activity showing higher TFEB cluster scores (Fig. 6.A-I). *GBA*-PD/DLB cases had lower total GCase activity thus showing higher amount of TFEB clustering compared to wild-type *GBA* cases (sPD/PDD and controls), which had lower cluster scores. Remarkably, the case (ID24) bearing three *GBA* variants (p.Asp140His, p.Glu326Lys, and p.Thr369Met) showed the lowest total GCase enzymatic activity and very high mean TFEB cluster score. No correlation between total GCase enzymatic activity and mean TFEB cluster score was observed in aSyn-positive cells (r=-0,215, p=n.s.) (Fig. 6.A-II). Similar correlation analysis between mean TFEB nuclear score and total SN GCase activity in aSyn-negative cells identified a modest positive correlation which was not significant (Spearman’s correlation, r=-0,213 p=n.s.) (Fig. S6.A-I).

**Figure 6:**
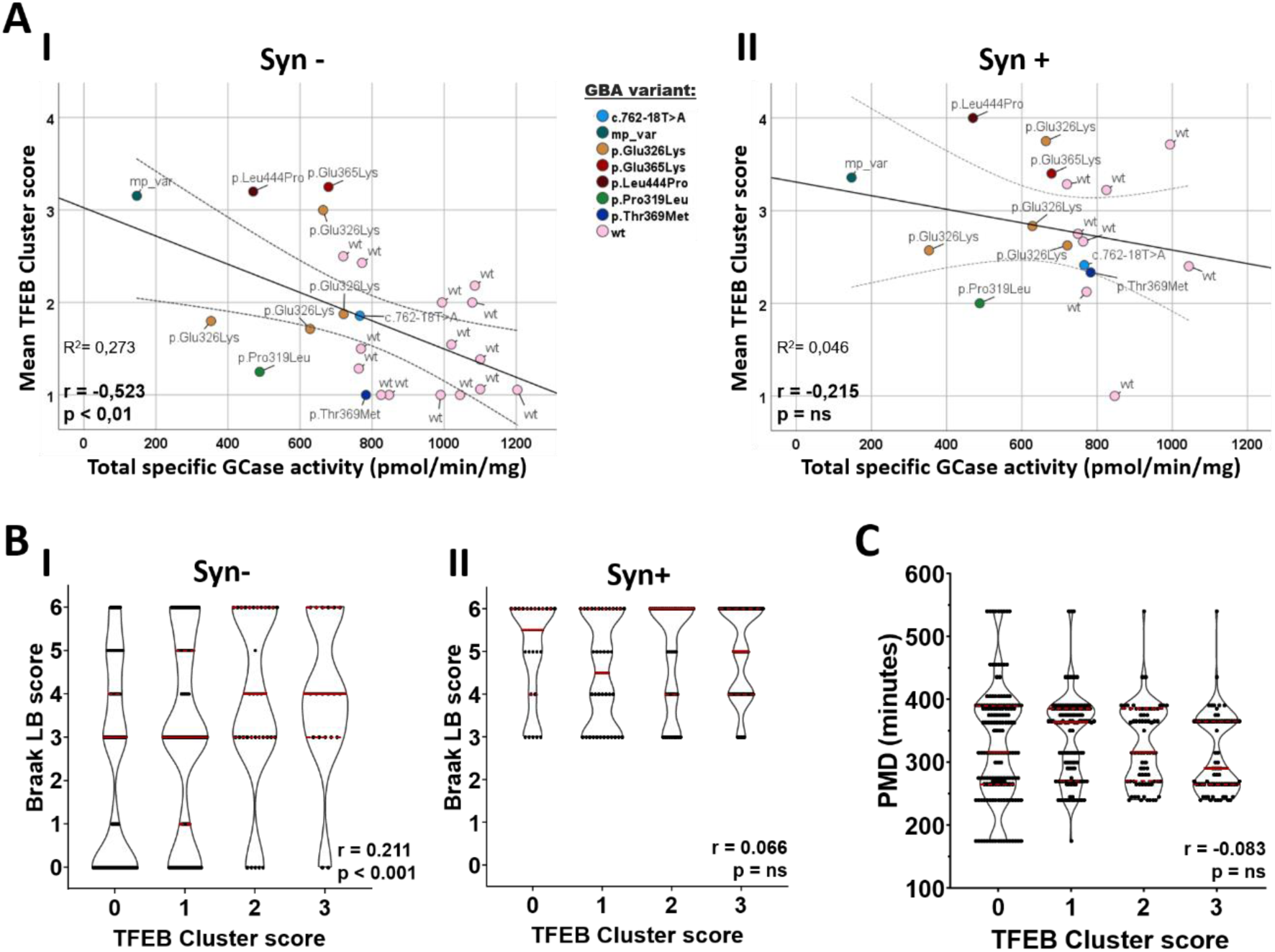
TFEB cluster score correlates with bulk tissue GCase enzymatic activity and with disease progression, as defined by Braak LB staging. **A:** Pearson’s correlation analysis between mean TFEB cluster score and total GCase enzymatic activity from bulk SN tissue (previously measured in [49]). Wild-type (wt) and GBA mutation-carrier cases are color-coded as indicated in the graph (mp_var = multiple variants). Dotted lines indicate mean 95% confidence interval. A negative correlation between the readouts is observed (r= -0.523, p< 0.01) in aSynuclein-negative (Syn-) cells (*A-I*). No significant correlation is observed in aSynuclein-positive (Syn+) cells (r= -0.215, p= ns) (*A-II*). **B:** Frequency distribution of TFEB cluster score and Braak LB disease stages, as a measure of disease progression. Spearman’s correlation analysis reveals a statistically-significant positive correlation between the scores when analyzing Syn-cells (B-*I*) (r = 0.211, p < 0.001). The association is not significant in Syn+ cells (*B-II*) (r = 0.066, p = ns). **C:** Frequency distribution of post mortem delay (PMD) and TFEB cluster score. Spearman’s correlation analysis reveals no statistically-significant correlation between PMD and TFEB cluster scores (r = -0.083, p = ns).

**Figure 7:**
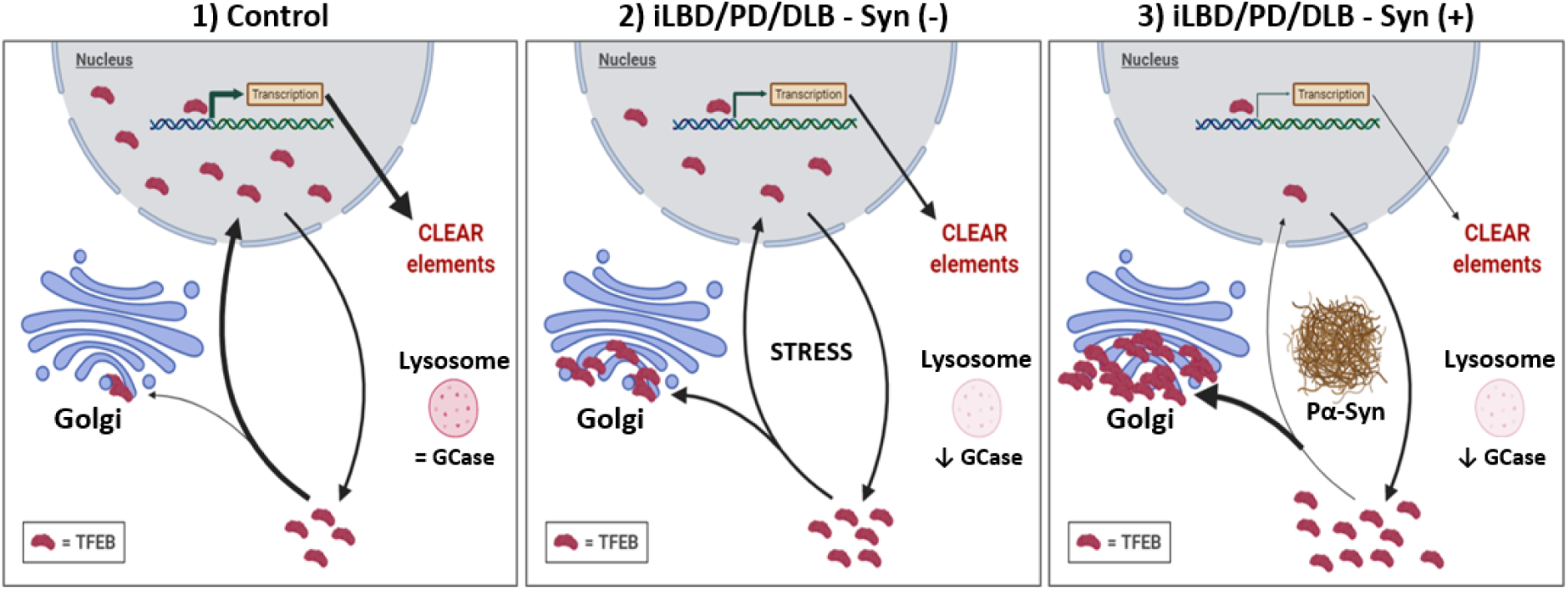
Hypothetical model of early TFEB redistribution and clustering upstream of pathological accumulation of pSer129 aSyn in PD/DLB neurons. In dopaminergic neurons from the SN of control cases, physiological localization of TFEB is preserved, which results in the physiological transcription of CLEAR network genes necessary for the correct function of the autophagy and lysosomal pathway (Panel 1). In PD/DLB and iLBD neurons, nuclear TFEB is reduced and TFEB clusters begin to form at the Golgi network in neurons prior apparent accumulation of pathological pSer129 aSyn (Pα-Syn), possibly in response to cellular stress triggered by a reduction in GCase activity and resulting lysosomal impairment. The observed TFEB redistribution is associated with an overall reduction of the transcription of CLEAR elements (Panel 2). Neurons with apparent aSyn cytopathology present more severe TFEB clustering at the TGN and a reduced nuclear pool (Panel 3) compared to PD/DLB neurons without intracellular aSyn (Panel 2). The observation that alterations of TFEB distribution occurs in aSyn (-) cells, both in GBA-related and sporadic PD/PDD cases, as well as in iLBD donors, indicates a role for TFEB in the cellular mechanisms leading to aSyn accumulation..

In order to study the association between the cluster score and disease progression, as defined by Braak LB staging, we analyzed the correlation between Braak LB score an TFEB cluster score (Fig. 6.B). Higher Braak LB score was associated with higher amount of TFEB clusters in aSyn-negative cells (r=0.211, p<0.001) (Fig. 6.B-I). The same effect was not observed when only analyzing aSyn-positive cells (r=0.066, p=n.s.) (Fig. 6.B-II). The observation that TFEB clustering associates with disease progression suggest its involvement in disease development. Doing so specifically in cells with have not yet developed aSyn cytopathology further indicate its involvement prior aSyn deposition.

To exclude the possibility that the observed TFEB clustering pattern could be influenced by differences in post-mortem delay (PMD), we analyzed the correlation between the TFEB score and PMD (Fig. 6.C). No significant correlation could be observed between the two outcomes thus confirming that the TFEB clustering measurement is not influenced by time differences between death and tissue collection (Spearman’s correlation, r=-0.083, p=n.s.). Similar analysis for nuclear TFEB has identified a modest but significant negative correlation between higher semi-quantitative scores and PMD (Spearman’s correlation, r=-0.105, p=0.014) (Fig. S6.C). This indicates that part of the observed effect of TFEB nuclear translocation by semi-quantitative scoring can be explained by differences in PMD between subjects. Nonetheless, no significant difference in PMD between the groups can be observed.

## Discussion

A growing number of pathological studies have focused on dissecting the determinants of ALP impairment in synucleinopathies. While several of these studies revealed mild effects on different ALP components in the context of PD/DLB, these results are sometimes contradictive [64, 65]. In the present study, we demonstrated that alterations in the subcellular localization of TFEB, master regulator of ALP, in nigral neuromelanin-containing neurons is strongly associated with PD and DLB in post-mortem human SN tissue. In line with earlier studies, we observed less nuclear localization of TFEB in nigral dopaminergic neurons of sPD/DLB and *GBA*-PD/DLB donors compared to age-matched control subjects, while increased cytoplasmic retention was associated with its clustering at the Golgi apparatus. Interestingly, these effects were also observed in prodromal PD/DLB (iLBD), indicating that altered TFEB localization initiates during the preclinical stage of PD/DLB. Moreover, TFEB deregulation was accompanied by an overall reduction in CLEAR genes mRNA expression levels and GCase enzymatic activity in bulk tissue, indicating that the TFEB impairment might result or be resulting from alterations of the ALP regulation. The effects were more pronounced in the *GBA*-PD/DLB group compared to sPD/DLB.

At the cellular level, TFEB clustering was notably increased in cells without detectable pSer129 aSyn cytopathology in *GBA* carriers and iLBD cases and was more severe in late pathological stages (as defined by Braak LB stages) and increased in neurons with intracellular pSer129 aSyn deposition. This finding suggests that TFEB impairment happens prior to pathological aSyn accumulation and worsens during the pathological advancement and progression of intracellular PD/DLB pathology (Fig. 6).

In 2013, Decressac et al. showed that adeno-associated virus (AAV) vector-mediated overexpression of human wild-type aSyn impairs TFEB in the midbrain [40], reflected by dynamic changes in CLEAR gene products (both mRNA expression levels and protein levels) over time, decreased relative protein expression levels of TFEB in nuclear vs cytoplasmic tissue fractions by Western blot [40]. In addition, decreased nuclear TFEB immunoreactivity was observed in sPD patients compared to controls.

In our study, we aimed to reproduce and expand on this latter finding in PD populations with and without *GBA*-mutations and in iLBD cases, using multiple labeling experiments and high-resolution microscopy allowing more insight into the detailed subcellular localization of TFEB and its relation to the presence subcellular pSer129 aSyn pathology. In line with the result of Decressac et al. [40], we observed significantly reduced nuclear TFEB immunoreactivity in dopaminergic neurons in the SNpc and an overall reduction of CLEAR genes expression levels in the midbrain of patients with sPD/DLB and, particularly, *GBA*-PD/DLB. The differences observed between the sporadic and *GBA*-related group can be ascribed to the existence of a more acute TFEB phenotype caused by mutations in *GBA*, which results in reduced GCase activity and has been previously associated with ALP impairment [30, 66-68]. Supporting this, the subcellular immunoreactivity patterns for two patients with severe *GBA* variants leading to very low residual GCase activities [49] – one donor carrying the pathogenic L444P variant and one donor with three *GBA* variants (p.Asp140His, p.Glu326Lys, and p.Thr369Met) – were characterized by extensive TFEB clusters and reduced nuclear labelling, occupying a substantial volume of the Golgi apparatus (Fig. 2,3) both in cells with and without aSyn pathology.

Apart from PD, reduced expression of TFEB in nuclear tissue fractions has been reported in different neurodegenerative diseases, including Alzheimer’s disease and Amyotrophic lateral sclerosis [69, 70]. These results demonstrate that nuclear TFEB localization is reduced in different neurodegenerative diseases in which protein aggregation takes place, thus suggesting that impaired nuclear translocation of TFEB is related to impaired protein homeostasis in neurons.

Besides neuronal cytoplasmic and nuclear TFEB punctae, we observed larger perinuclear immunopositive clusters co-localizing with Golgi markers. A link between TFEB and the Golgi apparatus was previously established in gene ontology analyses of the CLEAR network, showing an important role for TFEB in the transcriptional control elements of the Golgi apparatus [71]. Moreover, recent evidence has suggested a role of the Golgi apparatus in ALP functioning in early stages of autophagy, amongst others in mannose 6-phosphate receptors-mediated sorting of lysosomal enzymes [72], as a source of membranes of double-layered membranes for ALP components [73, 74], and for the formation of autophagosomes [75]. These findings together suggest the Golgi system might be a regulatory hub at the mTOR-TFEB axis.

Although accumulation of TFEB at the Golgi apparatus has not been reported before, a similar phenotype was described for ATG9A – an ALP component involved in early stages of autophagosome formation and intimately linked to the subcellular localization of TFEB [76]. In genetic mouse and cellular models of adaptor protein 4 (AP-4) deficiency and in human skin fibroblasts of patients with AP-4 mutations, accumulation of ATG9A at the Golgi was observed and was associated with axonopathy and an impaired ability to dispose cytoplasmic aggregates of huntingtin [77-79]. TFEB interacts with lysosomes via the interaction with its regulator mTORC1. Recent literature identified the existence of perinuclear clusters of lysosomes localized at the Golgi [80, 81], which allows for the activation of mTORC1 by a non-canonical pool of its activator protein, Rheb, which is localized at the Golgi. This mechanism has been shown to downregulate autophagy [82]. Hao *et al*. [80] demonstrated that the induction of cellular stress by starvation, which has been shown to lead to the activation of a TFEB-mediated response [35], promotes a dynein-dependent retrograde transport of lysosomes to the perinuclear Golgi complex. The existence of a pool of Rheb activator protein at the Golgi and the description of stress-induced association of lysosomes with the Golgi might explain the colocalization of the TFEB clusters with the Golgi identified in this study. Although future experiments are clearly needed to shed light on the causes and consequences of TFEB accumulation at the Golgi apparatus under diseased and healthy conditions, our observations highlight that the Golgi apparatus might be an important regulatory center for ALP function under conditions of cellular stress. In line with this, we observed signs of cellular stress, such as altered nuclear and cellular morphology and increased DNA condensation, in hESC-derived neurons displaying perinuclear TFEB clusters.

Previous studies demonstrated a high structural homology between aSyn and 14-3-3 proteins, a phospho-serine binder and a negative regulator of TFEB [61, 62, 83]. This family of proteins has been shown to regulate cellular response to stress and nutrient availability pathways [84]. This includes the inhibition of common negative regulators of mTORC1, such as TSC2 and PRAS40. aSyn has been shown to co-immunoprecipitate with 14-3-3 proteins and to interact to proteins which are common binders of, and regulated by, 14-3-3 proteins [62]. In the present study we observed limited colocalization of either total aSyn or pSe129-aSyn with TFEB or TFEB clusters. However, due to this structural homology between 14-3-3 proteins and phosphorylated aSyn, it cannot be excluded that phosphorylated aSyn might interact with binders of 14-3-3 proteins thereby negatively regulating mTORC1 and thus leading to TFEB hyperphosphorylation, its reduced nuclear translocation and reduced functionality. This might result in the initiation of alternative processes aimed at TFEB activation other than the one involving mTORC1-lysosome, such as one at the Golgi membrane level. Further investigation is needed to elucidate the mechanisms involved in this pathway.

Additionally, the observation of lysosomal interaction with existing clusters at the Golgi suggests that TFEB in the clusters might still be functionally active. This further advocates the possibility that the increased association of TFEB with the Golgi might be a physiological response to intracellular stress present prior intracellular aSyn accumulation. The observation of TFEB clusters in cells without intracellular aSyn deposition may indicate the existence of cellular stress prior to the manifestation of aSyn cytopathology and suggests the involvement of this process before aSyn accumulation. This proposition is supported by the existing correlation between disease progression, as quantified by the Braak staging, and TFEB clustering score in aSyn-negative and not in aSyn-positive cells as demonstrated by this study (Fig. 6). Furthermore, the observation that TFEB cluster formation negatively correlates with total GCase activity, especially in the more severe phenotypes such as the *GBA*-PD/PDD group, further suggest that cytoplasmic retention of TFEB could be triggered by lysosomal dysfunction. Therefore, elucidating the functional meaning and the mechanisms involved in the translocation of TFEB at the Golgi might give further insights on the processes involved in early disease progression and lead to the identification of a potential early therapeutic window.

In conclusion, our results on TFEB subcellular localization in postmortem human brain tissue have demonstrated a link between TFEB deregulation and the development of *GBA*-related and sPD/DLB and may suggest its involvement in the molecular disease development prior to pathological pSer129 aSyn accumulation. Our observations support an important role of TFEB in the pathogenesis of *GBA* carriers and sPD/DLB, thereby confirming the potential of targeting the TFEB pathway as a possible approach for urgently-required disease-modifying therapies for synucleinopathies.

## Acknowledgements

We thank dr. Marialuisa Quadri (Erasmus MC, University Medical Center Rotterdam, Department of Clinical Genetics) for the support with the genetic analysis of *GBA* genotyping and dr. Tommaso Beccari (University of Perugia, Department of Pharmaceutical Sciences) for the input and support with the enzymatic activity assays on post-mortem human brain. We thank Bram van der Gaag (Amsterdam UMC, Vrije University Medical Center, department of Anatomy and Neurosciences, Amsterdam) for the help with image acquisition. We also thank Daniel Mona, dr. Markus Britschgi (Roche Pharma Research and Early Development, Neuroscience and Rare Diseases Discovery and Translational Area, Roche Innovation Center Basel) and dr. Wagner M. Zago (Prothena Biosciences Ltd., Dublin, Ireland) for providing the primary antibody against pSer129 aSyn and its direct-labelling with fluorophores.

## Ethical approval and consent to participate

Post-mortem human brain tissue from clinically diagnosed and neuropathologically-verified donors with PD, DLB, iLBD as well as non-demented controls was collected by the Netherlands Brain Bank (NBB, www.brainbank.nl). In compliance with all local ethical and legal guidelines, informed consent for brain autopsy and the use of brain tissue and clinical information for scientific research was given by either the donor or the next of kin. The Code of Conduct and Ethical Declaration of the NBB are publicly accessible [85-87]. The procedures of the Netherlands Brain Bank (Amsterdam, The Netherlands) were approved by the Institutional Review Board and Medical Ethical Board (METC) from the VU University Medical Center (VUmc), Amsterdam.

## Competing interests

The authors declare no competing interests. M.L.M, T.E.M., V.U. and R.J. are or were full-time employees of Roche/F. Hoffmann-La Roche Ltd, and they may additionally hold Roche stock/stock options.

