## Supplementary information for "Altered TFEB subcellular localization in nigral dopaminergic neurons of subjects with prodromal, sporadic and *GBA*-related Parkinson’s disease and Dementia with Lewy bodies"

Tim E Moors[1,2] & Martino L Morella[1,2], Cesc Bertran-Cobo[1,2], Hanneke Geut[1,2], Vinod Udayar[3], Evelien Timmermans-Huisman[1,2], Angela MT Ingrassia[1,2], John JP Brevé[1,2], John GJM Bol[1,2], Vincenzo Bonifati[4], Ravi Jagasia[3], Wilma DJ van de Berg[1,2,†]

**Affiliations:**

[1] Amsterdam UMC, location Vrije University Medical Center, section Clinical Neuroanatomy and Biobanking, department of Anatomy and Neurosciences, Amsterdam, The Netherlands.

[2] Amsterdam Neuroscience, program Neurodegeneration, Amsterdam, The Netherlands

[3] Roche Pharma Research and Early Development; Neuroscience and Rare Diseases Discovery and Translational Area; Roche Innovation Center Basel.

[4] Erasmus MC, University Medical Center Rotterdam, Department of Clinical Genetics, the Netherlands.

† Corresponding author:
Wilma D.J. van de Berg, PhD
Dept. of Anatomy & Neurosciences, chair section Clinical Neuroanatomy and Biobanking
Amsterdam Neuroscience
Amsterdam UMC, location VU University Medical Center
O2 building, room 13 E11
De Boelelaan 1108
1081 HZ Amsterdam, The Netherlands

### Supplementary Figures and Tables

**Table S1: Donors selected for this study.** More detailed demographics were provided in Moors et al 2017 [1]

| ID (ID in [1]) | Pathological diagnosis | Age | Sex | GBA variant | Severe GBA variant (Y/N) | Braak LB stage | Braak NFT score | CERAD amyloid score | IHC exp. | GCase activity exp. | qPCR Exp. | Thal phase |
|---|---|---|---|---|---|---|---|---|---|---|---|---|
| **1 (1)** | control | 77 | F | - | - | 0 | 1 | B | x | x | - | 2 |
| **2 (2)** | control | 84 | F | - | - | 0 | 1 | O | - | - | x | 0 |
| **4 (4)** | control | 79 | F | - | - | 0 | 1 | O | - | - | x | 0 |
| **5 (5)** | control | 83 | F | - | - | 0 | 1 | B | - | x | - | 2 |
| **6 (6)** | control | 76 | F | - | - | 0 | 2 | O | x | x | x | 0 |
| **7 (7)** | control | 76 | M | n.a. | n.a. | 0 | 0 | O | - | - | x | 1 |
| **8 (8)** | control | 70 | F | - | - | 0 | 2 | A | x | x | x | 1 |
| **9 (9)** | control | 83 | F | - | - | 0 | 2 | B | x | - | - | n.a. |
| **10 (10)** | control | 70 | F | - | - | 0 | 2 | A | x | x | - | 2 |
| **12 (12)** | control | 78 | F | - | - | 0 | 2 | A | x | x | - | 2 |
| **14 (14)** | control | 79 | M | - | - | 0 | 2 | A | - | - | x | 3 |
| **15 (15)** | control | 83 | M | - | - | 1 | 1 | A | x | x | x | 1 |
| **17 (17)** | PD | 81 | F | - | - | 5 | 0 | O | x | x | - | 0 |
| **19 (19)** | PD | 74 | M | - | - | 5 | 1 | O | x | x | - | 0 |
| **22 (22)** | PD | 81 | M | - | - | 5 | 1 | O | x | x | x | 0 |
| **23 (23)** | PD | 80 | M | - | - | 6 | 2 | B | - | - | x | 2 |
| **25 (25)** | PD | 72 | M | - | - | 6 | 1 | A | x | x | - | 3 |
| **27 (27)** | PD | 78 | M | - | - | 4 | 0 | A | x | x | x | n.a. |
| **28 (28)** | PD | 73 | M | - | - | 4 | 3 | O | - | - | x | 0 |
| **29 (29)** | PD | 81 | F | - | - | 6 | 3 | A | - | - | x | n.a. |
| **30 (30)** | PD | 76 | M | - | - | 6 | 2 | O | - | - | x | 0 |
| **34 (34)** | DLB | 70 | M | - | - | 6 | 1 | A | - | - | x | 3 |
| **35 (35)** | DLB | 83 | M | - | - | 5 | 2 | B | - | - | x | n.a. |
| **36 (36)** | DLB | 78 | M | - | - | 6 | 2 | C | - | - | x | 2 |
| **39 (39)** | DLB | 75 | M | - | - | 6 | 2 | B | x | x | - | 3 |
| **41 (41)** | DLB | 78 | F | - | - | 6 | 2 | B | x | x | x | n.a. |
| **44 (44)** | DLB | 83 | M | - | - | 6 | 0 | O | x | x | x | 0 |
| **45 (45)** | DLB | 74 | M | - | - | 4 | 1 | B | x | x | x | 2 |
| **16 (16)** | PD | 80 | F | c.762-18T>A | n.a. | 6 | 1 | C | x | x | - | 1 |
| **18** | PD | 84 | M | p.Pro319Leu | n.a. | 5 | 1 | A | x | x | - | 2 |

| | | | | | | | | | | | | |
|---|---|---|---|---|---|---|---|---|---|---|---|---|
| **(18)** | | | | | | | | | | | | |
| **21 (21)** | PD | 76 | M | p.Glu326Lys | N | 6 | 1 | O | x | x | x | 0 |
| **24 (24)** | PD | 83 | F | p.Asp140His, p.Glu326Lys, and p.Thr369Met | Y | 4 | 1 | O | x | x | x | 0 |
| **26 (26)** | PD | 65 | M | p.Leu444Pro | Y | 6 | 1 | B | x | x | x | 3 |
| **37 (37)** | DLB | 72 | M | p.Glu326Lys | N | 6 | 1 | O | x | x | - | 0 |
| **38 (38)** | DLB | 80 | F | p.Glu326Lys | N | 5 | 1 | O | x | x | x | 0 |
| **42 (42)** | DLB | 81 | F | p.Thr369Met | N | 4 | 1 | A | x | x | x | 2 |
| **43 (43)** | DLB | 69 | M | p.Glu326Lys | N | 6 | 3 | B | x | x | - | 4 |
| **50** | iLBD | 93 | F | - | - | 3 | 2 | O | x | - | - | 0 |
| **51** | iLBD | 86 | F | - | - | 3 | 3 | A | x | - | - | 1 |
| **52** | iLBD | 98 | M | - | - | 3 | 2 | O | x | - | - | 0 |

**Table S2:** Details of the antibodies used in the present study.

| Reference/ Art no. | Source | Epitope | Host (clonality) | Dilution (µg/ml) |
|---|---|---|---|---|
| 11A5 | Prothena | P–Ser129 α–syn | Mouse (monoclonal) | IHC: 1:20000 (0,3)<br>- |
| sc-7012 (N-19) | Santa Cruz | aSyn (aa 1-60) | Goat (polyclonal) | IHC: 1:100 (1)<br>- |
| A303-673A | Bethyl | TFEB | Rabbit (polyclonal) | IHC: 1:100 (10)<br>WB: 1:1000 (1) |
| ab220695 | Abcam | TFEB | Rabbit (polyclonal) | IHC: 1:25 (n.a.)<br>WB: 1:100 (n.a.) |
| HPA049532 | Atlas Antibodies | TFEB | Rabbit (polyclonal) | -<br>WB: 1:170 (15,26) |
| MBS120432 | MyBioSource | TFEB | Mouse (monoclonal) | -<br>WB: 1:1000 (0,1) |
| ab270614 | Abcam | TFEB | Rabbit (monoclonal) | -<br>WB: 1:1000 (0,98) |
| #86843 | Cell Signaling Technology | Phospho-TFEB (Ser122) | Rabbit (polyclonal) | -<br>WB: 1:1000 (0,005) |
| ab2636 | Abcam | TFEB | Goat (polyclonal) | -<br>WB: 1:1000 (0,5) |
| NBP1-49643 | Novus Biologicals | TGN46 | Rabbit (polyclonal) | IHC: 1:150 (6,67)<br>- |
| HPA021230 | Atlas Antibodies | GOLGA2 | Rabbit (polyclonal) | IHC: 1:100 (7)<br>- |

| | | | | |
|---|---|---|---|---|
| H4B4 | Developmental Studies Hybridoma Bank | LAMP-2 | Mouse (monoclonal) | IHC: 1:200 (n.a.) - |
| MAB422 | EMD Millipore | Cathepsin D | Mouse (monoclonal) | IHC: 1:100 (10) - |

**Table S3:** Number of nigral neuromelanin-containing dopaminergic neurons with and without pathology scanned in the different diagnostic groups for the semi-quantitative analysis of TFEB

| | | **Pathology** | | |
|---|---|---|---|---|
| | | **α-SYN (–)** | **α-SYN (+)** | **Total** |
| **Diagnosis** | **Control** | 98 | 0 | 98 |
| | **iLBD** | 74 | 31 | 105 |
| | **sPD/DLB** | 54 | 55 | 109 |
| | ***GBA*-PD/DLB** | 57 | 72 | 129 |
| | **Total** | 283 | 158 | **441** |

**Table S4:** List of primer used in the qPCR experiments

| Target | Primer sequence<br>5' 3' | UPL probe number | Holding stage (95°C) | Denaturation (95°C) | Annealing (T(°C)/sec) | Extension (65°C) |
|---|---|---|---|---|---|---|
| *HEXA* | F: ACACTTCCGCTGTGAATTGCTG<br>R: GCTCCACTACCATTCACCTACA | # 15 | 15 min | 15 sec | 59 °C/30 sec | 40 sec |
| *MAP1LC3A* | F: GACCATGTCAACATGAGCGAG<br>R: CGTAGACCATATAGAGGAAGCC | # 15 | 15 nmin | 15 sec | 59 °C/30 sec | 45 sec |
| *SOSTM1* | F: GAAGCTGCCTTGTACCCACA<br>R: CCGATGTCATAGTTCTTGGTCTG | # 16 | 15 min | 15 sec | 60 °C/30 sec | 55 sec |
| *UGCG* | F: CAAAGCGATAGCTGACCGAG<br>R: CTGGCAACAAAGCATTCTGAAATTG | # 4 | 15 min | 15 sec | 59 °C/30 sec | 45 sec |
| *VPS35* | F: CTCTTCTGGTCTGGCAGAAAC<br>R: TCTTCTCGAATCTTTTGGATAAGCTG | # 52 | 15 min | 15 sec | 60°C/30 sec | 55 sec |
| *GALC* | F: GTGTGTTCATTGCAGGAAGAG<br>R: GTAACTTCAACACGTCCTAAAGC | # 6 | 15 min | 15 sec | 59 °C/30 sec | 45 sec |
| *GBA* | F: CCTACTCATGCTGGATGACCA<br>R: CCAATGTACAGCAATGCCATGAA | # 43 | 15 min | 15 sec | 58 °C/35 sec | 30 sec |
| *TFEB* | F: GAGATGACCAACAAGCAGCTC<br>R: AGCTCAGCCATGTTCATGCC | # 17 | 15 min | 15 sec | 56 °C/30 sec | 30 sec |
| *OAZ1* | F: CACCATGCCGCTCCTAAG<br>R: ACAGCAGTGGAGGGAGAC | # 74 | 15 min | 15 sec | 57°C/30 sec | 30 sec |
| *POL2RF* | F: CATGTCAGACAACGAGGACAA<br>R: TCCAAGTCATCTAGCCCTTCA | # 25 | 15 min | 15 sec | 60°C/30 sec | 30 sec |
| *POLR2A* | F: CGCATCATGAACAGCGATGAG<br>R: GCAGGGTCATATCTGTCAGCAT | # 69 | 15 min | 15 sec | 59°C/30 sec | 35 sec |
| *PES1* | F: AGATGCAGAGGCTGGTTCA<br>R: CTCACTCTCCTCCTCCT | # 21 | 15 min | 15 sec | 59°C/30 sec | 30 sec |

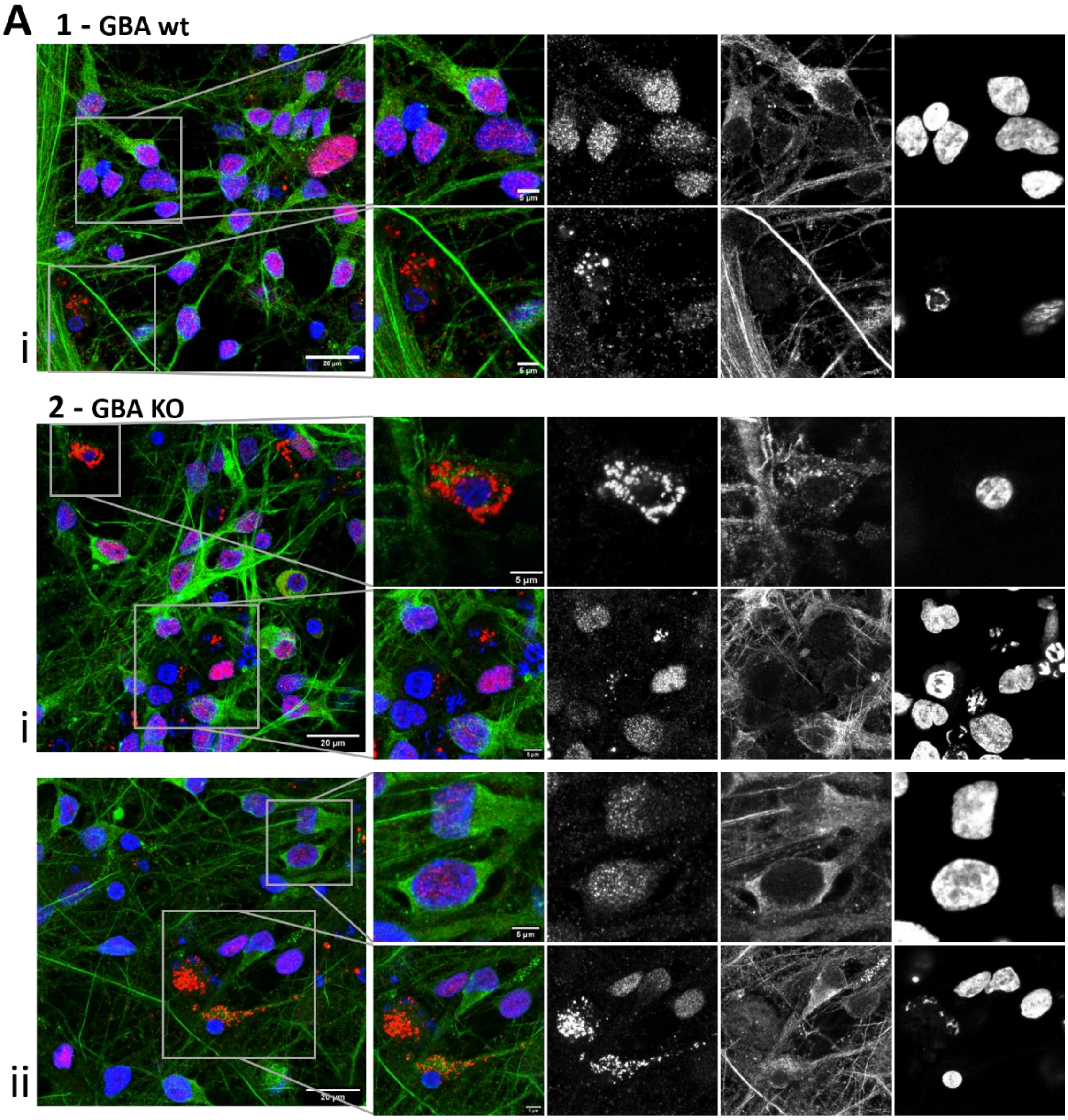
A
1 - GBA wt
i
20 µm
5 µm
5 µm
2 - GBA KO
i
20 µm
5 µm
ii
20 µm
5 µm

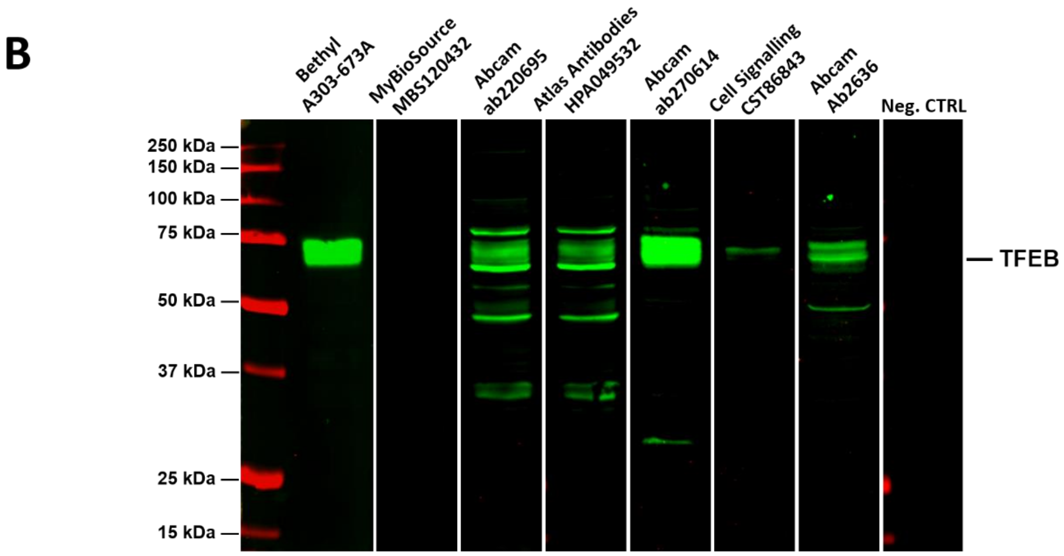
B
Bethyl A303-673A
MyBioSource MBS120432
Abcam ab220695
Atlas Antibodies HPA049532
Abcam ab270614
Cell Signalling CST86843
Abcam Ab2636
Neg. CTRL
250 kDa
150 kDa
100 kDa
75 kDa
50 kDa
37 kDa
25 kDa
15 kDa
TFEB

### Figure S1: Specificity of TFEB antibodies.

**A:** TFEB clusters are observed in human embryonic stem cell-derived neurons in vitro with TFEB A303-673A antibody. Representative raw confocal overview images (left panels) and zoom-ins (right panels) of hESC-derived isogenic dopaminergic neurons stained for TFEB (red) and MAP2 (green) showing TFEB perinuclear clusters similar of what observed human *post-mortem* brain. Cells displaying TFEB clusters were often displaying signs of cellular stress, such as altered cellular morphology, reduced nuclear shape and DNA condensation. *1* - GBA wild-type (wt) neurons. *2* - GBA knock-out (KO) neurons. No difference in the percentage of cluster-positive neurons could be observed among the two lines. Scale bar = 20/5 µm as indicated in each panel. **B:** Comparison of specificity of TFEB antibodies by western blot. Comparison of the specificity of several commercially-available antibodies for TFEB by SDS as indicated.

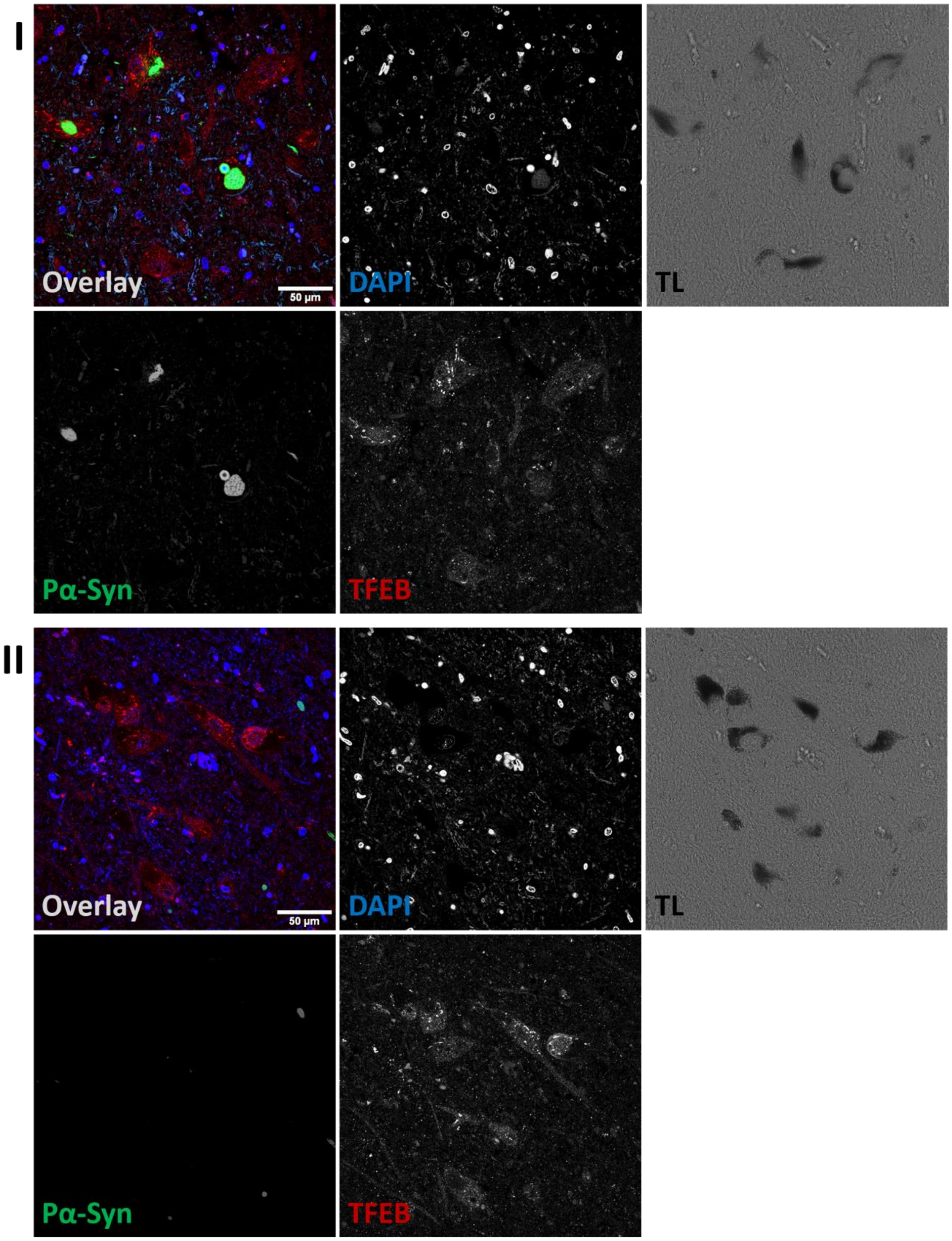


**Figure S2: Representative low magnification fluorescent IHC images of TFEB clusters with Bethyl A303-673A antibody.**

Representative transmitted light (TL, right panels) and raw CLSM (other panels) images of SNpc from donor ID19 (sPD, *I*), ID50 (iLBD, *II*). Staining for TFEB (red) and Ser129-phosphorylated aSyn (Pα-Syn, green) reveals the presence of intracellular aSyn cytopathology and perinuclear TFEB clusters. Scale bar = 50 µm.

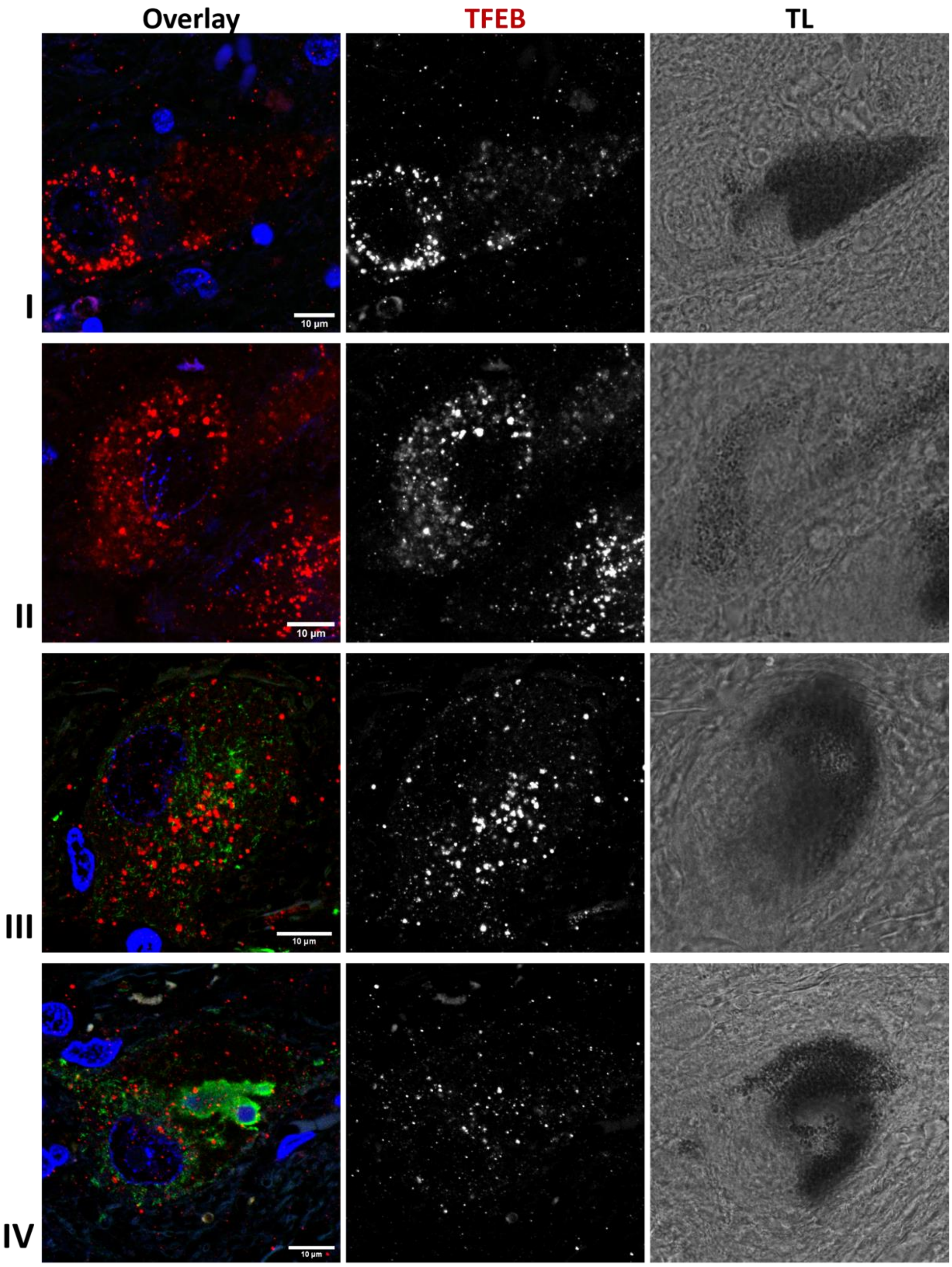


**Figure S3: Representative fluorescent IHC images of TFEB immunopositive clusters with Abcam ab220695 antibody.**

Representative transmitted light (TL, right panels) and raw CLSM (other panels) images of SNpc from donor ID17 (sPD, *I*), ID39 (sDLB, *II-III*), ID25 (sPD, *IV*) stained for TFEB (red) and Ser129-phosphorylated aSyn (green) showing the presence of perinuclear TFEB positive clusters immunoreactive to Abcam ab220695 TFEB antibody. Scale bar = 10 µm.

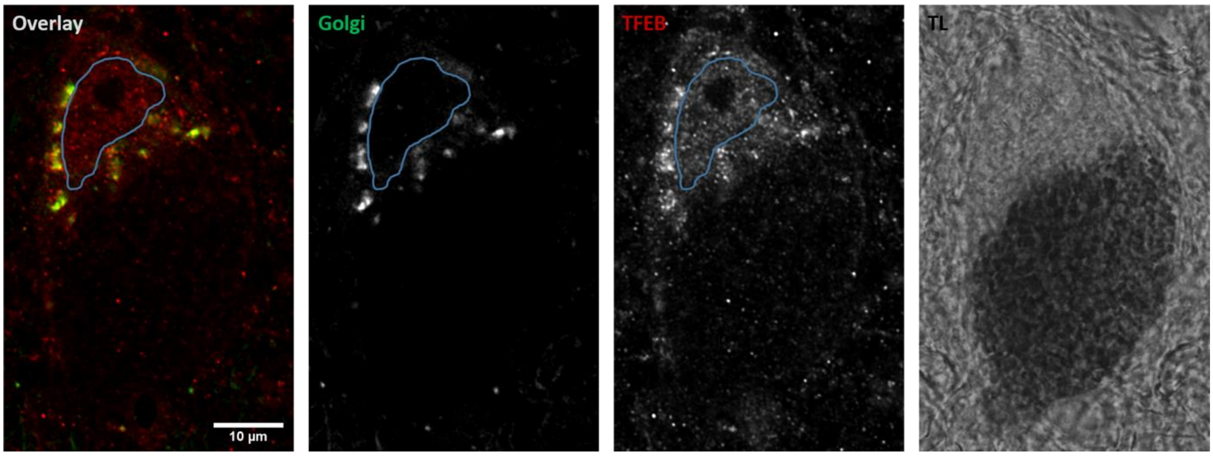


**Figure S4: Representative sequential fluorescent IHC image of TFEB immunopositive clusters localized at the Golgi.**

Representative transmitted light (TL, right panel) and raw CLSM (other panels) images of SNpc from donor stained for TFEB (red) and the cis-Golgi marker GOLGA2 (green) stained with sequential fluorescent IHC as described in the methods sections showing the localization of perinuclear TFEB clusters at the Golgi. Scale bar = 10 µm.

.

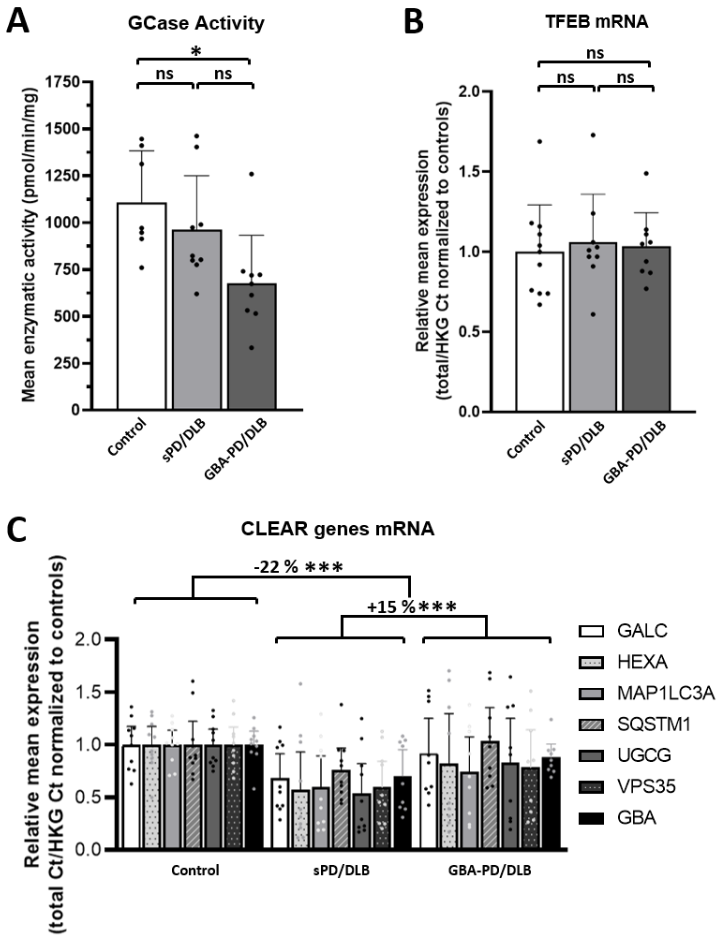


**Figure S5: GCase activity and selected CLEAR genes expression are decreased in the medial frontal cortex of sPD-DLB and GBA-PD/DLB patients.**

**A:** Mean total GCase enzymatic activity quantification in bulk SN tissue from sPD/DLB, GBA-PD/DLB patients and controls expressed as pmol/min/mg of total protein. Mean ± SD; N≥7/group, n=3. **B, C:** mRNA quantification by qPCR in sPD/DLB, GBA-PD/DLB patients and controls calculated as total Ct normalized on HKGs and expressed as fold-change compared to the control group. **B:** Quantification of TFEB mRNA. Mean ± SD; N≥5/group, n=3. **C:** CLEAR genes mRNA quantification. Mean ± SD. N≥5/group, n=3. HKG: housekeeping genes; GALC: galactosylceramidase; HEXA: hexosaminidase subunit alpha; GBA: ß-glucocerebrosidase; MAP1LC3A: microtubule associated protein 1 light chain 3; SQSTM1: sequestosome 1; UGCG: UDP-glucose ceramide glucosyltransferase; VSP35: retromer complex component. *p < 0.05; ***p < 0.001; † p < 0.05 vs Control.

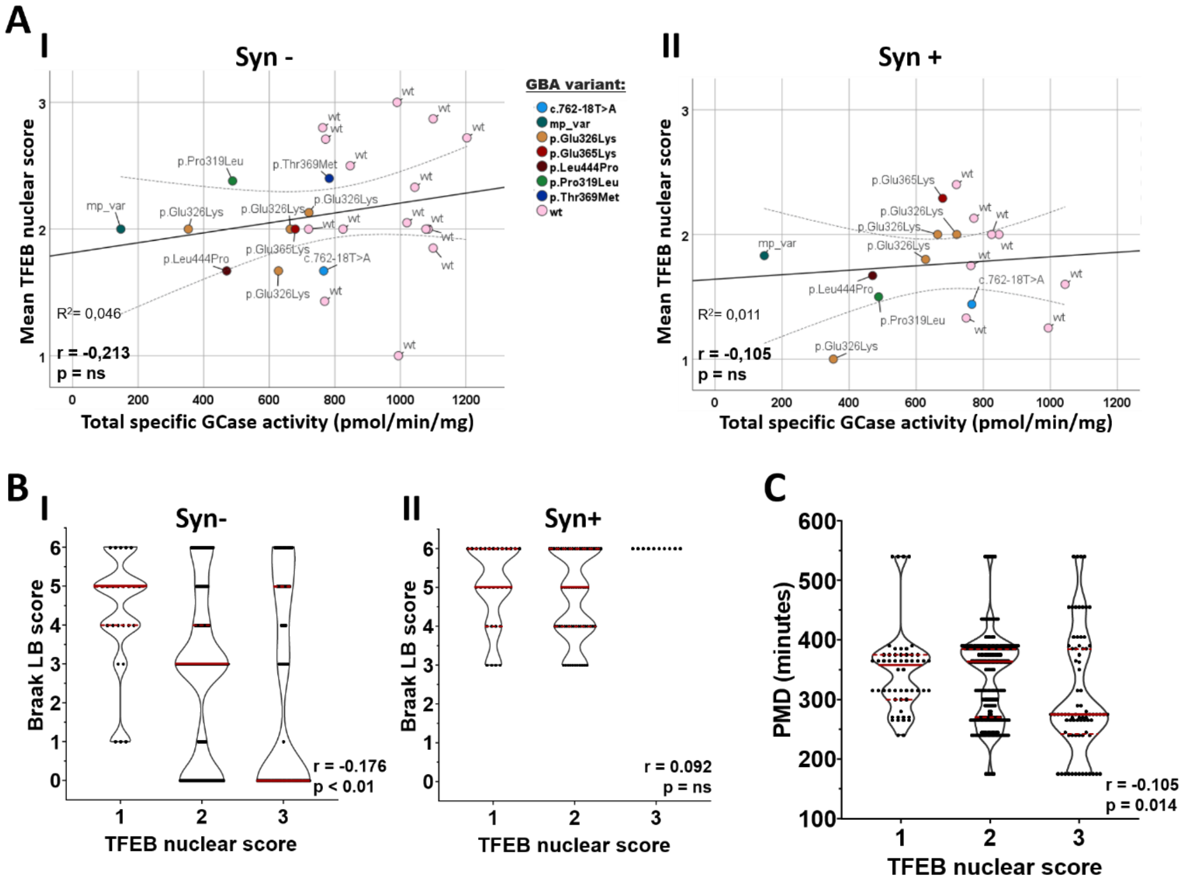


**Figure S6: TFEB nuclear score correlates with bulk tissue GCase enzymatic activity and disease progression, as defined by Braak LB staging.** **A:** Pearson's correlation analysis between mean TFEB cluster score and total GCase enzymatic activity from bulk SN tissue (previously measured in [1]). Wild-type (wt) and GBA mutation-carrier cases are color-coded as indicated in the graph (mp_var = multiple variants). Dotted lines indicate mean 95% confidence interval. No statistically-significant correlation between the readouts is observed both in aSynuclein-negative (Syn-) cells (r= -0.213, p= ns) (*A-I*) and in aSynuclein-positive (Syn+) cells (r= -0.105, p= ns) (*A-II*). **B:** Frequency distribution of TFEB cluster score and Braak LB disease stages, as a measure of disease progression. Spearman's correlation analysis reveals a statistically-significant positive correlation between the scores when analyzing Syn- cells (*B-I*) (r = -0.176, p < 0.01). The association is not significant in Syn+ cells (*B-II*) (r = 0.092, p = ns). **B:** Frequency distribution of post mortem delay (PMD) and TFEB cluster score. Spearman's correlation analysis reveals no statistically-significant correlation between PMD and TFEB cluster scores (r = -0.105, p = 0.014).

**Supplementary Material and Methods**

*Western Blotting*

Total cell lysate from human monocyte cell line HP1 was prepared in SDS sample buffer (final: 50 mM Tris-HCl, 10% glycerol, 2% SDS, 1% 2-mercaptoethanol, 12.5 mM EDTA, 0.02 % bromophenol blue) containing 50 mM DTT and then heated for 10 minutes at 95°C. Samples containing 40 µg of total protein were loaded per each well of a 10% acrylamide (29:1) TRIS/Glycine SDS gels (1.5 mm/10-well) and run at 150V for 1-1.5 hr. Proteins where than transferred on nitrocellulose membrane in Transfer Buffer (25mM Tris, 200mM glycine) containing 10% methanol. After blotting, the membranes were briefly washed in TBS-T (50 mM Tris, 150 mM NaCl, pH 7.6 + 0.1% Tween 20) and blocked for 30 min at RT in Odyssey blocking buffer:TBS (1:1)(Li-Cor). Blots were then cut in vertical strips and each was incubated with different TFEB antibodies diluted in Blocking buffer (Odissey Bocking buffer:TBS=1:1) or in blocking buffer alone (negative controls) and incubated overnight at 4°C. The following TFEB antibodies were used (see Table S2 for details): TFEB A303-673A (Bethyl), TFEB ab220695 (Abcam), TFEB HPA049532 (Atlas Antibodies), TFEB MBS120432 (MyBioSource), TFEB ab270614 (Abcam), Phospho-TFEB (Ser122) #86843 (Cell Signaling Technology), TFEB ab2636 (Abcam). Dilutions were not optimized for the assay and are based on pre-existing literature. After primary, blots where washed in TBS-T and incubated with secondary antibodies (Goat anti-Rabbit, IDRye800CW, 1:10000, Li-cor 926-32211 or Donkey anti-Mouse IRDye800CW 1:10000, Li-cor 926-32212) diluted in Blocking buffer for 1 hr at RT. After washing with TBS-T and rinsing once in TBS, the strips were imaged together using Odyssey SA scanner and Image Sudio Software (Li-Cor).

*Sequential multiplex IHC*

Post-mortem FFPE brain section were processed as described in the *Immunohistochemical staining procedure* section of Materials and methods until and including the antigen retrieval step. After, the slides were quenched in 3% hydrogen peroxide in TBS for 30 minutes at RT and then washed three times for 5 minutes in TBS and incubated in Blocking buffer (3% normal goat serum in TBS) for 30 minutes at RT. The sections were than incubated with TFEB A303-673A (Rabbit, 1:100, Bethyl) primary antibody in Blocking buffer over-night at 4°C. In the morning, the sections were washed three times in TBS and incubated with Envision HRP anti-Rabbit (DAKO #K4003) for 1 hr at RT. After washing twice with TBS and once with 1M Tris-HCl, the slides were incubated with Alexa Fluor 594 Tyramide Reagent (Invitrogen #B40957) for 15 mins at RT and immediately washed 2 times in 1M Tris-HCl and once in TBS. In order to remove the antibodies from the tissue sections (stripping) and proceed with a sequential multiplex staining, the tissue sections were processed for antigen retrieval as previously described. The slides were than incubated with an anti-GOLGA2 primary antibody (Rabbit, 1:400 , HPA021230, Atlas Antibodies) in Blocking buffer for 4 hr at RT and then incubated with a directly-labelled secondary antibody (Goat anti-Rabbit Alexa Fluor™ 647, 1:400, Thermo Fisher #A-21245) in Blocking buffer containing 1 µg/ml fo DAPI for 2 hr at RT. After washing, sections were mounted in Mowiol mounting solution using glass cover slips (Art. No.: 630-2746; Glaswarenfabrik Karl Hecht, Sondheim, Germany).

*Differentiation of human embryonic stem cells to neurons*

-Human embryonic stem cells

Human embryonic stem cells (hESCs) were obtained with ethical approval under Swiss research registry (BAG-hES-IMP-0031) from the UK Stem Cell Bank (UKSCB) (Steering comm. appl. SCSC07-14). Cell line source: Human Embryonic Stem Cell Line SA001, Cellartis AB; NIH Human Embryonic Stem Cell Registry no. 0085; origin info: male, ethnicity and age N/A. *GBA* KO was generated from the same hESC line as previously reported (GBA1_hESC_GBA#1−/−_10.18.11; identifier: CEBe033-A-1) [2].

-Neural precursor cells

Neural precursor cells (NPCs) were generated from human embryonic stem cells (hESC) according to a previously published protocol [3]. In brief, hESCs were plated as single cell suspension in AggreWell-800 (STEMCELL Technologies, #34815) to generate embryoid bodies (EBs) and maintained in NPC1 media (DMEM/F-12-Neurobasal mix medium [1:1 mix of DMEM/F-12 medium (supplemented with 1%

GlutaMax, Thermo Fisher #31331093) and neurobasal medium (Thermo Fisher #21103049; supplemented with 2% B27 supplement, 2% N2 supplement, and 50 µM 2-mercaptoethanol (ThermoFisher #12587010, #17502048, #31350010 respectively] supplemented with 5 ng/ml FGF-2, 250 ng/ml noggin (R&D Systems or Peprotech) supplemented with 20 µM SB 431542 (Tocris #1614) for 5 days. EBs were than plated in plates coated with polyornithine-laminin (PL) and maintained in the same medium for 3 day to form neural rosettes. At day 4, neural rosettes were manually dissociated and plated in PL-coated plates in NPC1. After reaching confluence, cells were dissociated and plated at 100 000 cells/$cm^2$ on PL-coated plates in NPC2 media (DMEM/F-12-Neurobasal mix medium supplemented with 10 ng/ml FGF-2 (Peprotech), 10 ng/ml EGF (R&D Technologies), and 20 ng/ml BDNF (Peprotech). Cells were then passaged every 2-3 days for about 15 times with a gradual decrease in cell density to 25 000 cells/$cm^2$ over the first 10 passages and maintained in NPC2 media with daily media changes.

-Differentiation to neuronal cells

Differentiation of NPCs to neurons was perform according to a previously-published protocol [4]. NPCs were plated at 10 000 cells/ $cm^2$ in PL-coated flasks and cultured for one week in DMEM/F-12-Neurobasal mix medium supplemented with 100 ng/ml FGF-8 (Peprotech), 200 ng/ml sonic hedgehog (Peprotech), and 100 µM ascorbic acid 2-phosphate (Sigma). Cells were then re-plated at 50 000 cells/$cm^2$ in neurobasal medium supplemented with 20 ng/ml BDNF, 10 ng/ml glial cell-derived neurotrophic factor (GDNF; Peprotech), 500 µM dibutyryl cyclic AMP (Sigma), and 100 µM ascorbic acid 2-phosphate and differentiated for 42 days before assays.
